# Biofilm initiation associated with cell orientation patterning in *Escherichia coli*

**DOI:** 10.64898/2026.03.26.714369

**Authors:** Fumiaki Yokoyama, Kazumasa A. Takeuchi

**Affiliations:** Department of Physics, The University of Tokyo, Bunkyo-ku 113-0033, Tokyo, Japan; Microbial Lifeform Dynamics Group, Okinawa Institute of Science and Technology, Onna-son, Kunigami-gun 904-0495, Okinawa, Japan; Universal Biology Institute, The University of Tokyo, Bunkyo-ku 113-0033, Tokyo, Japan; Institute for Physics of Intelligence, The University of Tokyo, Bunkyo-ku 113-0033, Tokyo, Japan

**Keywords:** biofilm, Rcs pathway, microfluidics, active nematics, topological defects

## Abstract

Unicellular microorganisms can transition to multicellular states that enhance survival under environmental fluctuations. In bacteria, one such state is the biofilm, defined by the production of an extracellular matrix. Although many aspects of biofilm formation have been extensively studied, how matrix production is spatially initiated within a growing population remains largely unknown. Here we show that cell orientation patterning is associated with the spatial initiation of Rcs stress-response activation, which is known to be involved in extracellular matrix production. Using an *Escherichia coli* strain reporting mechanically induced stress responses, we observed that Rcs activation was accompanied by mechanically regulated out-of-plane growth under agar-pad confinement. Notably, this activation preferentially emerged near dense topological defects, where cell orientation mismatches and growth-induced pressure builds up. By controlling confinement geometry with microfluidic devices, we regulated cell orientation patterns and defect distributions, thereby biasing the spatial pattern of Rcs activation. These findings indicate an association between cell orientation patterning and the spatial activation of a stress-response pathway involved in extracellular matrix production, suggesting a possible physical link to biofilm initiation.

**Significance:** Biofilms help bacteria survive environmental change, but it remains unclear how specific locations within a growing population begin to adopt biofilm-related behaviors. We show that the collective alignment of *Escherichia coli* cells influences where the Rcs stress-response pathway, which regulates extracellular matrix production, becomes activated. Activation occurred preferentially in regions where the orderly alignment of neighboring cells broke down and where growth-generated pressure is known to build up. By changing the geometry of the space confining the population, we altered cell alignment and shifted the spatial pattern of Rcs activation. These findings reveal an association between the physical organization of a bacterial population and local stress-responsive gene regulation, suggesting a possible link to the spatial initiation of biofilm development.

## 1 Introduction

Unicellular microbial organisms often make a transition to multicellular states that help them to survive under environmental fluctuations [1]. In bacteria, biofilms constitute one such multicellular life form, which is composed of dwelling cells and extracellular matrix by definition [2]. According to this definition of biofilm, when a cell population starts producing extracellular matrix, it turns into a biofilm. Nevertheless, the mechanism underlying the initial step of this biofilm transition process remains poorly understood, in particular regarding where and how the matrix production is initiated. While cells inside biofilm tend to grow more slowly than the periphery, cells before turning into biofilm grow actively and push each other in optimal conditions [3]. Considering the bacterial mechanotransduction, we hypothesized that mechanical responses caused by cell contact upregulated gene expression of mechano-inducible extracellular matrix production, triggering biofilm initiation.

Bacterial mechanical responses induce gene expressions through surface molecules and subsequent signaling systems, as shown for various bacterial species [4, 5]. These mechanically upregulated genes encode various types of molecules that can function as extracellular matrices [4, 6–8]. Among them, colanic acid (CA) is one of the representative extracellular matrices of *Escherichia coli* [9] and induced through the Rcs (regulator of capsule synthesis) stress-response pathway as a mechanical response of cell deformation, imposed by surface contact [10] and spatial confinement [11]. This CA matrix is an extracellular heteropolymer of four types of saccharides [12] and works as a support for large three-dimensional (3D) biofilm structure [13] and a barrier against antibiotics [14] and bacteriophages [11], showing its significant contribution to biofilm formation and collective survival. Along the Rcs pathway, *rprA* is simultaneously upregulated with CA production through mechanotransduction [15]. An *rprA*-based fluorescent transcriptional reporter has therefore been used to visualize mechanically induced Rcs activation in individual *E. coli* cells under confinement [11]. The Rcs activation can in turn promote CA production through regulation of the CA biosynthetic genes.

In dense populations of rod-shaped bacteria, mechanical responses arise heterogeneously because growth-induced pressure builds up non-uniformly through collective cell alignment and cell–cell collisions [16]. In species such as *E. coli* [17], *Pseudomonas aeruginosa* [18], *Myxococcus xanthus* [19, 20], and *Alcanivorax borkumensis* [21], the spatial distribution of pressure within a population can be inferred from patterns of cell orientation. Because rod-shaped cells tend to align along one another through mechanical contact, growing bacterial monolayers form ordered orientation patterns with topological defects, which refer to singular points in the cell orientation field. At the defects, the local cell orientation becomes undefined. When cells grow around these defects, growth pressure builds up, causing cells to buckle [22–30]. As a result, orientation patterns and their associated topological defects organize the spatial distribution of pressure and drive collective bacterial behaviors [27, 31–34].

Despite extensive studies of biofilms, a central unresolved question is how spatially localized extracellular matrix production is initiated within a growing bacterial population. Here, we addressed this question by introducing cell orientation patterns as a framework linking collective cell organization to activation of a matrix production-involving pathway in bacterial monolayers. Using confined *E. coli* populations, we found that high Rcs activation was associated with regions influenced by dense topological defects. By controlling confinement geometry, we regulated cell orientation patterns, thereby localizing topological defects and biasing the spatial distribution of Rcs activation. Together, these results reveal an association between cell orientation patterning and the spatial activation of a stress-response pathway involved in matrix production, suggesting that collective cell organization may contribute to the physical processes underlying biofilm initiation.

## 2 Results

### 2.1 Activation of Rcs stress-response pathway by protruding cells from monolayers

To test Rcs activation during 3D biofilm formation, we observed growth and Rcs activation of *E. coli* cells using fast-maturating monomeric super-folder green fluorescence protein (msfGFP) [35] under the native promoter of *rprA*. We observed cells on a glass substrate under a soft agar pad with lysogeny broth (LB) at 37 °C by confocal microscopy (Fig. 1a), comparing two different focal planes: the bottom and 3 *µ*m higher (Fig. 1b sketch). Cells grew on the surface of the glass substrate two-dimensionally (2D) first and, once they covered almost the whole surface in the field of view, some cells within the colony protruded into the upper layer by invading the soft agar pad (Figs. 1b,c, Supplementary Videos 1 and 2). Comparison of the time series of the protruding area and Rcs activation shows that cells protruded into the upper layer first, then started Rcs activation approximately 20 min later (Fig. 1d). Although some cells in the bottom layer of the colony also showed Rcs activation (Fig. 1b at 40 and 60 min), the Rcs activation per area, quantified at the bottom images, in the protruding region was higher than that in the non-protruding region (Fig. 1e). This Rcs activation in the protruding area remained approximately four- to sixfold higher than in the non-protruding area over 10–60 min during Rcs activation (Fig. 1f). These results suggest that cells in a growing 2D colony protruded and activated the Rcs pathway, most likely followed by production of CA as an extracellular matrix component through the Rcs mechanical response pathway, which senses cell–cell and cell–substrate contacts as well as cell deformation.

**Fig. 1.**
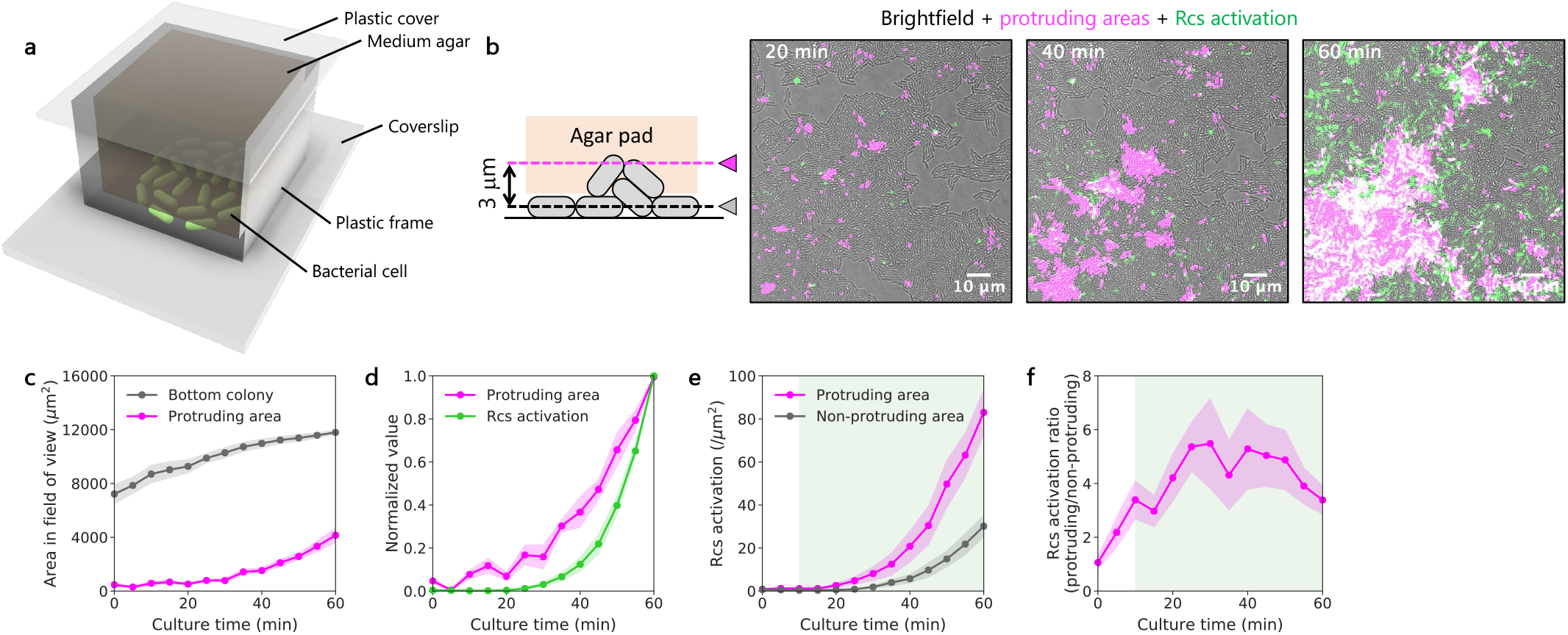
Activation of Rcs stress-response pathway within cell-protruding areas. **a**, A sketch of experimental setup using an agar pad. Bacterial cells on a glass substrate are covered with an agar pad of LB. A plastic cover and a 3D-printed frame are put on an agar pad and adhered by double-sided tape to a glass coverslip to prevent evaporation from the agar. **b**, Time evolution of cell protrusion from bottom layer of colony, and Rcs activation. A side view of the agar pad set-up with two focal planes at the bottom and 3 *µ*m higher is drawn (left). Time-lapse overlays of cell protrusion from the bottom layer of a colony (magenta) and Rcs activation visualized by msfGFP expression (green) are shown (right). The sketches in the panels **a** and **b** are not to scale. **c**, Time evolution of areas occupied by the bottom layers of the colonies, and protruding cells in field of view. **d**, Time evolution of the protruding area (magenta) and the msfGFP fluorescence intensity (green) reporting Rcs activation, both normalized by the values at 60 min. **e**, Time evolution of Rcs activation per area in the protruding and non-protruding regions. **f**, Time evolution of the ratio of Rcs activation per area in the protruding region, compared to that in the non-protruding region. The green shades in **e, f** show the time range of Rcs activation from 10 to 60 min. Each line plot shows the mean with standard errors (10 positions across 2 devices were used).

### 2.2 Characterization of topological defects in monolayer confiner devices

Mechanically driven responses in cell populations arise not only from global compression but also from localized pressure. In mammalian cell monolayers, biological responses such as apoptosis have been reported to occur at topological defects, where mechanotransduction is triggered by characteristic local mechanical force fields [36]. This defect-driven mechanotransduction motivated us to examine whether mechanically induced biological features, such as extracellular matrix production, are similarly associated with cell orientation patterns and topological defects in bacteria.

Cell orientational fields are dynamically reorganized by collisions between neighboring microcolonies. This makes it difficult to relate cell orientation to Rcs activation, which is induced with a temporal delay (Fig. 1d). Therefore, we developed a microfluidic device, which we call monolayer confiner, to confine bacterial cells into a monolayer and let them grow within the confinement over time by supplying culture medium (Fig. 2a, Extended Data Fig. 1), referring to the previously reported design [37].

**Fig. 2.**
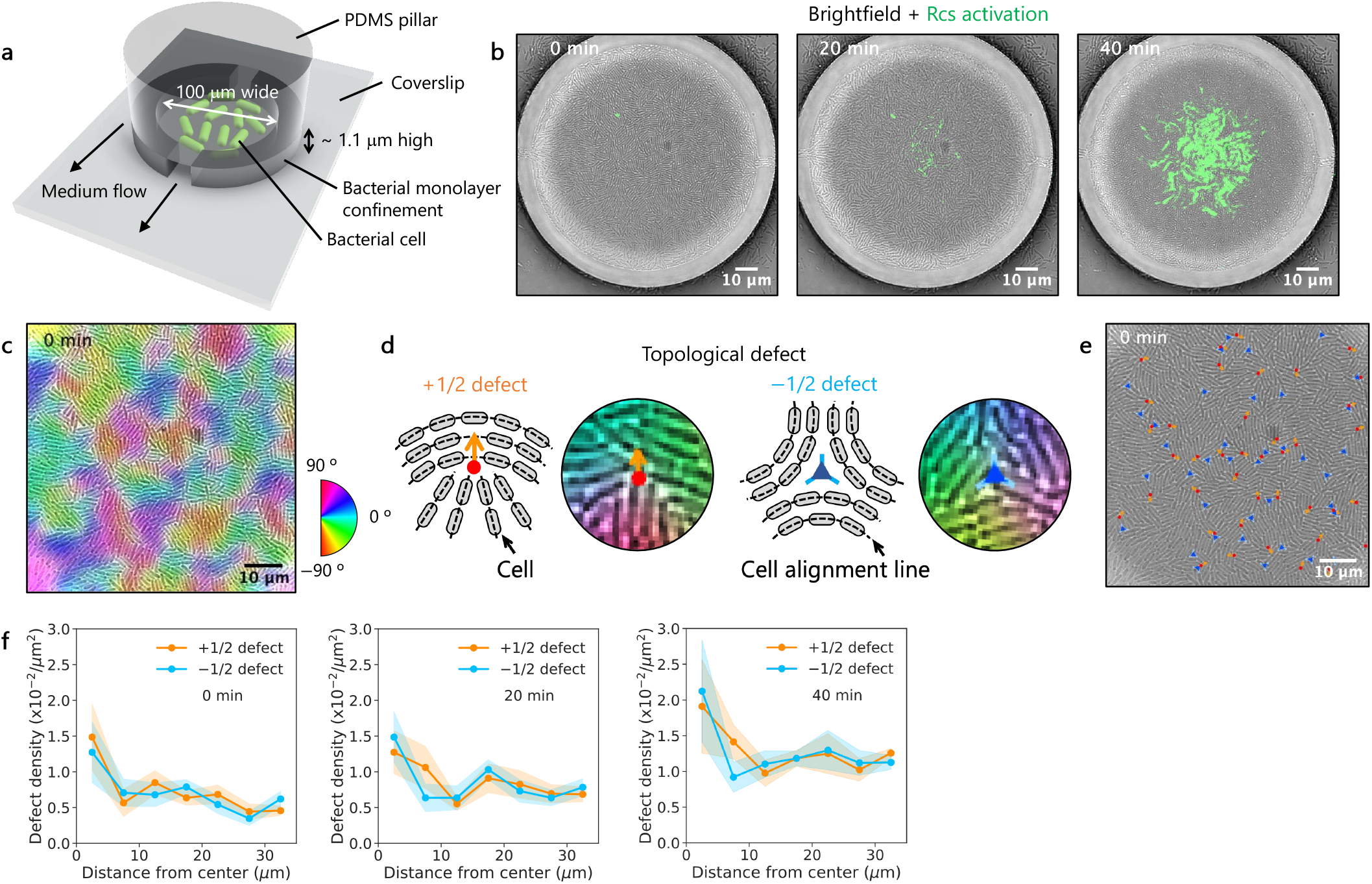
Characterization of topological defects in monolayer confiner devices. **a**, A sketch of experimental setup using microfluidic device (see also Extended Data Fig. 1). Bacterial cells within a chamber beneath a pillar of a polydimethylsiloxane (PDMS) device on a glass coverslip are shown. The chamber has a pair of inlet and outlet to let fresh medium come in for cell culture. Its height is approximately 1.1 *µ*m to form a monolayer of *E. coli* cells (Extended Data Fig. 1). This sketch is not to scale. **b**, Time-lapse overlays of brightfield and msfGFP fluorescence images of cells growing and activating the Rcs pathway in a monolayer confiner device. The green color indicates fluorescence intensity of msfGFP associated with Rcs activation. **c**, A representative brightfield image of cells in a monolayer confiner device with orientational color codes. **d**, Sketches and images of *±* 1/2 topological defects. The centers of +1/2 and −1/2 topological defects are shown by a red circle and a blue triangle, respectively. The cells are colored depending on their orientation, indicated by the same color code as **c. e**, A representative brightfield image of cells in a monolayer chamber device with *±* 1/2 topological defects. **f**, Density of *±* 1/2 topological defects as a function of the distance from the colony center at 0, 20, 40 min (from left to right). Each line plot shows the mean with standard errors (6 positions across 2 devices were used).

For cell growth analysis in the chamber, we used images of cells that remained attached to the chamber under flow. (Supplementary Video 3). Cells in this monolayer confiner chamber grew and activated the Rcs pathway (Extended Data Fig. 2a,b). The colony area increased exponentially with time (Extended Data Fig. 2c–e). The distribution of Rcs activation expanded approximately isotropically, indicating that flow directions did not affect Rcs activation (Extended Data Fig. 2f,g). This implies that this matrix production is not controlled by diffusive small molecules, such as quorum-sensing autoinducers and nutrients, which typically follow flow directions [38].

For cell orientation analysis, we used images of chambers fully occupied by growing cells (Fig. 2b, Supplementary Video 4). Cell orientation was quantified using the structure tensor method [39, 40] (Fig. 2c, Extended Data Fig. 3a,b, Supplementary Video 5) [25, 41]. Using this orientation field, we detected topological defects. In the literature, two primary types of topological defects are known, referred to as +1/2 and −1/2 defects[42]. +1/2 defects exhibit a distinct comet-like shape, where cells fan out from a defect center indicated by the red circle in Fig. 2d (left). This creates anisotropic force fields, which result in defect locomotion along the defect axis indicated by the orange arrow in Fig. 2d (left) in extensile systems. In contrast, −1/2 defects form trefoil-like patterns around the blue triangle in Fig. 2d (right), producing more isotropic force fields. The differences in local force distribution between these defects have biological implications in bacteria [17–21]. +1/2 defects, with their active motility and cell growth, can serve as nucleation points for the formation of a secondary layer in bacterial colonies because of increased pressure [17, 19, 20]. Meanwhile, −1/2 defects tend to repel cells and open holes in a bacterial swarm [19] unless bacteria vertically tilt and 3D effects arise in growing population [17].

Defect organization was position-dependent within the colony. Cells in the chamber spontaneously generated nearly equal numbers of +1/2 and −1/2 topological defects (Fig. 2e,f, Extended Data Fig. 3c,d, Supplementary Video 6), and the densities of both defect types increased over time (Extended Data Fig. 3d). Throughout the observation period, defects were consistently more concentrated near the colony center than at the edge (Fig. 2f, Extended Data Fig. 3e). These results show that defect distributions were spatially non-uniform within monolayer confiner chambers, with distinct organization of +1/2 and −1/2 defects.

### 2.3 Activation of Rcs stress-response pathway around high density of topological defects

To investigate the contribution of *±* 1/2 topological defects to Rcs activation during biofilm initiation, we compared msfGFP fluorescence intensity associated with Rcs activation with defect positions in the monolayer-confining chambers. Time-lapse overlays of msfGFP fluorescence showed that Rcs activation was first initiated near the chamber center (Fig. 3a, Supplementary Video 7). A kymograph confirmed that Rcs activation began at the colony center at approximately 20 min and expanded outward to encompass the entire colony by 40 min (Fig. 3b). Although Rcs activation is initiated near the colony center in the monolayer confiner chambers, its spatial distribution within the central region is heterogeneous and exhibits a non-trivial pattern rather than a uniform or isotropic profile. This suggests that factors beyond colony geometry contribute to Rcs activation. We examined whether cell orientation and topological defects contribute to the observed spatial patterns of Rcs activation.

**Fig. 3.**
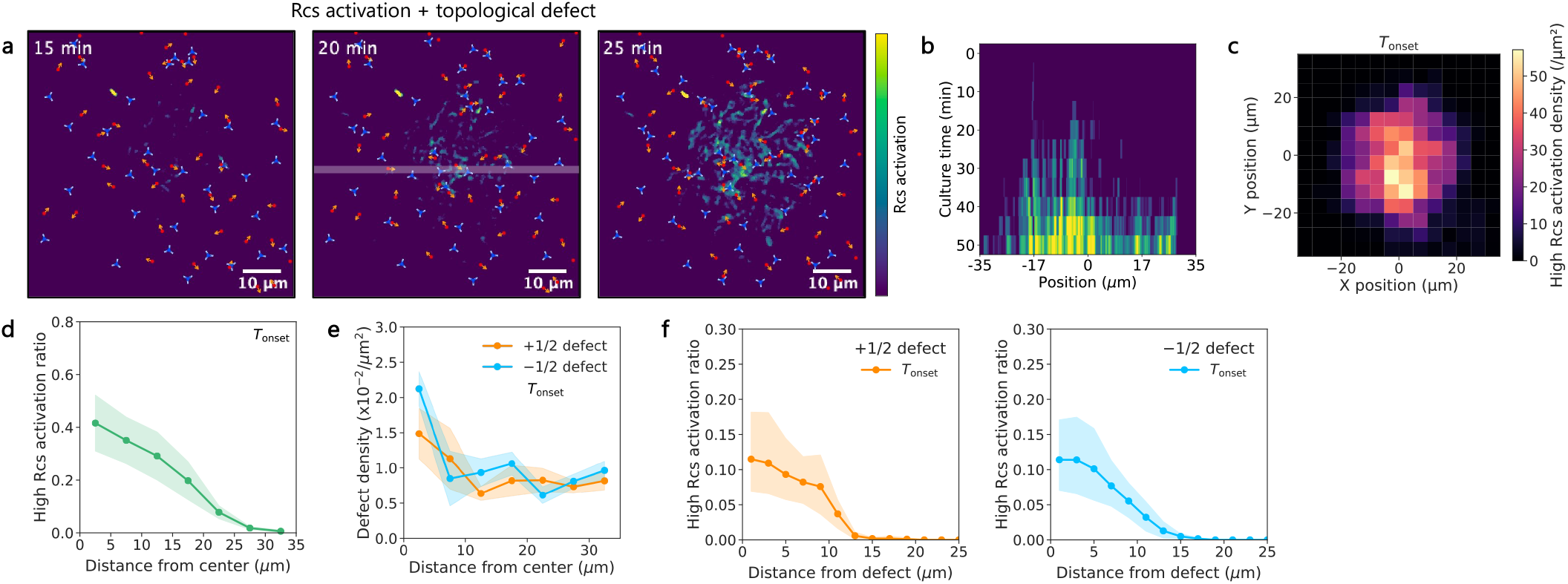
Activation of Rcs stress-response pathway around high density of topological defects. **a**, Time-lapse overlays of msfGFP fluorescence reporting Rcs activation and *±* 1/2 topological defects in a monolayer confiner device. The msfGFP fluorescence intensity reporting Rcs activation is shown using the viridis color scale. The red circles and blue triangles show +1/2 and −1/2 topological defects, respectively. The white line at 20 min shows the position for creating the kymograph in **b** along it. **b**, A kymograph of Rcs activation in a monolayer confiner device. The viridis color scale indicates the msfGFP fluorescence intensity reporting Rcs activation. **c**, 2D distribution of high Rcs activation pixel densities over colony at *T*_onset_. High Rcs activation pixels were defined as those with msfGFP fluorescence intensities exceeding the maximum value observed in the initial state, when cells fully occupied the chamber. **d**, 1D distribution of high CA pixel ratios as a function of the distance from the colony center at *T*_onset_. The ratio of high CA pixels was calculated by dividing the number of high Rcs activation pixels by the total pixel number. **e**, Density of *±*1/2 topological defects as a function of the distance from the colony center at *T*_onset_. **f**, High Rcs activation pixel ratios as a function of the distance from +1/2 and −1/2 topological defects (left and right, respectively) at *T*_onset_. The ratio of high Rcs activation pixels was calculated by dividing the number of high Rcs activation pixels by the total number of pixels at a given distance from defect (see Extended Data Fig. 5d). Each line plot shows the mean with standard errors (6 positions across 2 devices were used).

We defined the onset of Rcs activation as the time point at which the maximum msfGFP fluorescence increased by at least 20 units relative to the initial value (denoted by *T*_onset_), exceeding background levels after chamber filling and marking the initiation of biofilm formation. Colonies reached this threshold at a median time of 20 min (Extended Data Fig. 4). At this onset time, high Rcs activation pixels, whose intensities exceeded the maximum value observed when cells fully occupied the chamber, appeared near the colony center (Fig. 3c, Extended Data Fig. 5a).

To compare distributions of Rcs activation across monolayer confiner chambers, the numbers of high Rcs activation pixels at *T*_onset_ (and at *±* 5 min) were normalized by the total pixel number in radial bins to obtain high Rcs activation ratios. These ratios were elevated near the colony center (Fig. 3d, Extended Data Fig. 5b), coinciding with a high local density of *±* 1/2 topological defects (Fig. 3e, Extended Data Fig. 5c). Furthermore, the ratio of high Rcs activation pixels was calculated by dividing the number of high Rcs activation pixels by the total number of pixels at a given distance from defect (Extended Data Fig. 5d). This high Rcs activation ratio increased near the centers of *±* 1/2 topological defects and decreased with distance from the defects (Fig. 3f, Extended Data Fig. 5e,f). This Rcs activation was not due to a global upregulation of gene expression in centrally located cells, as tdTomato fluorescence driven by a constitutive promoter remained uniformly distributed across the colony (Extended Data Fig. 6). Together, these results show that both Rcs activation and topological defect density are elevated near the colony center in confined monolayers, implying a correlation between the two. However, this correlation alone does not exclude the possibility that their association arises from their shared central localization. To disentangle these effects, we next examined whether Rcs activation is specifically associated with the positions of topological defects rather than with colony center itself.

### 2.4 Localizing Rcs stress-response activation by regulating cell orientation

We next tested whether Rcs activation sites can be controlled by manipulating cell orientation and the positions of topological defects. To this end, we designed a microfluidic device containing square chambers of various sizes that confine bacteria into monolayers, which we called square confiners and channel-shaped compartments for comparison (Extended Data Fig. 7a,b). Because rod-shaped *E. coli* cells tend to align along chamber boundaries [43–45], chamber geometry provides a means to control orientational order and defect positions.

Square confiner chambers induced specific orientation fields and defect organization that depend on the confiner size. In smaller chambers with diagonal lengths of 10 and 20 *µ*m, cells exhibited ordered cell alignment with peaks of cell orientation around 0°, whereas in larger chambers with diagonal lengths of 30 and 40 *µ*m the orientation field became increasingly disordered (Fig. 4a,b, Supplementary Videos 8– 11). In chambers with diagonal lengths of 20 *µ*m, cell orientation patterns were classified as splay or bend at time points when cells fully occupied square chambers (Fig. 4c, Extended Data Fig. 8a). The splay configuration was associated with the persistent localization of *±* 1/2 defects near chamber corners (Extended Data Fig. 8b).

**Fig. 4.**
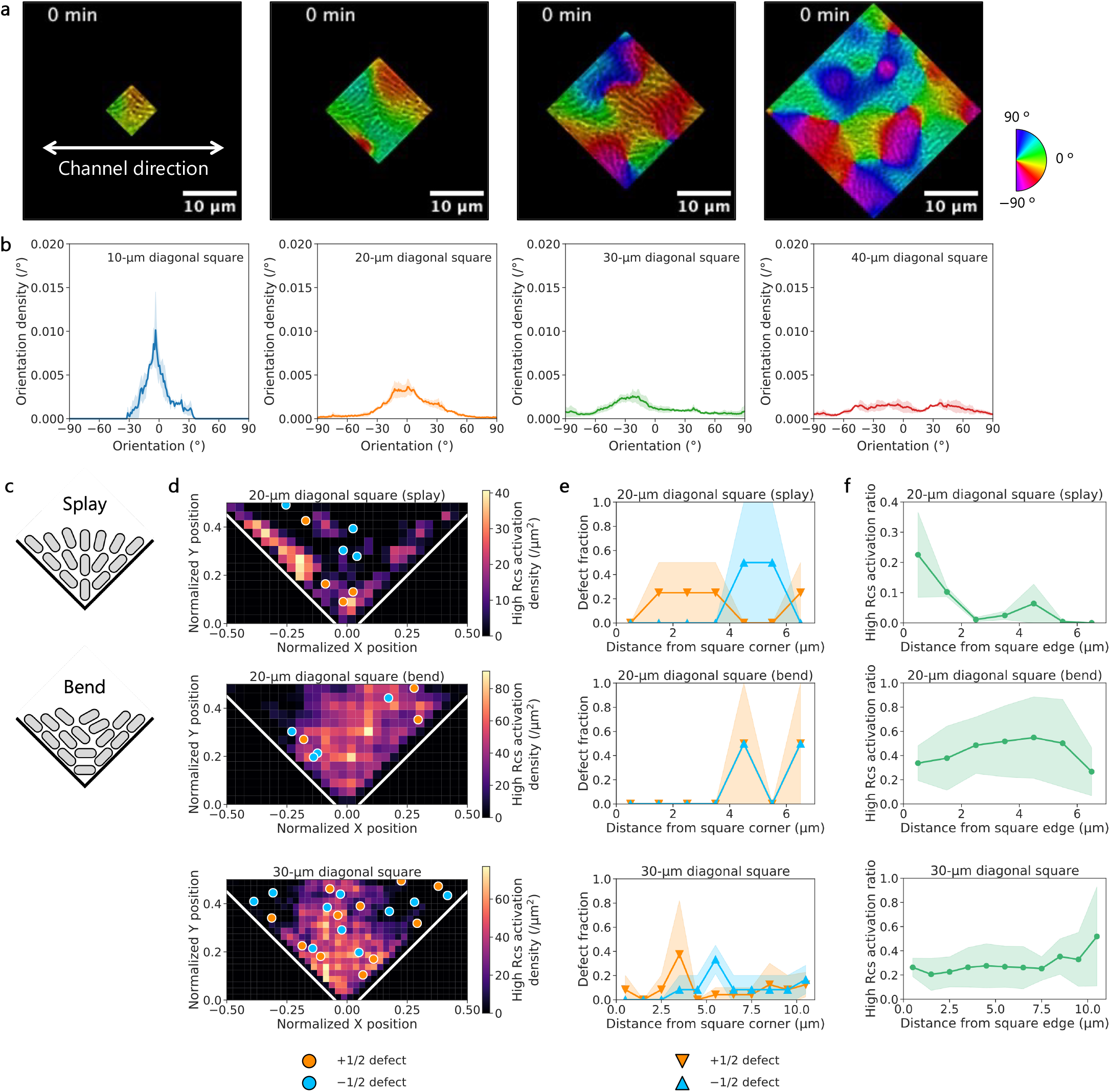
Localizing Rcs stress-response activation by controlling cell orientation. **a**, Representative cropped brightfield images of cells in square confiner devices with orientational color codes at *T*_onset_. Channel inlets/outlets were extended as indicated by the white arrow from the square chamber (see Extended Data Fig. 7). **b**, Count density of cell orientation in the square confiner devices *±* 20 min after chambers filled with cells. **c**, Sketches of splay (top) and bend (bottom) patterns of cell orientations. These sketches are not to scale. **d**, Spatial distribution of representative topological defects and high Rcs activation pixel densities in square confiner devices with diagonals of 20 and 30 *µ*m at *T*_onset_. Cell orientation patterns were classified into splay and bend at time points when cells fully occupied square chambers (Extended Data Fig. 8a). X and Y positions are normalized by the diagonal length of the square chambers. White lines delineate the edges of the square chambers. **e**, Mean number fraction of *±*1/2 topological defects as a function of the distance from the corner in square confiner devices with diagonals of 20 and 30 *µ*m at *T*_onset_. **f**, Distribution of high Rcs activation pixel ratios as a function of the distance from the edge in square confiner devices with diagonals of 20 and 30 *µ*m at *T*_onset_. Each line plot shows the mean with standard errors (≥ 4 square corners and ≥ 2 positions across ≥ 2 devices were used).

We measured Rcs activation and found a chamber-size-dependent spatial pattern associated with the characteristic cell orientation at *T*_onset_. In smaller chambers exhibiting splay cell orientation, the +1/2 defects preferentially localized near corner regions, resulting in a clear boundary-associated bias in +1/2 defect distribution, while −1/2 defects were distributed away from the corners (Fig. 4d,e, top). Under these conditions, high Rcs activation regions were enriched near the chamber boundary and decreased toward the interior (Fig. 4d,f, top). In contrast, in bend configurations, *±* 1/2 defects were predominantly found away from the chamber corners (Fig. 4d,e, middle), and Rcs activation extended across the chamber (Fig. 4d,f, middle). In smaller chambers with mixed cell orientation, *±* 1/2 defects showed no pronounced spatial localization, and Rcs activation was similarly delocalized across the chamber (Extended Data Fig. 8c–e). In larger chambers, defect distributions became more spatially distributed compared to smaller chambers (Fig. 4d,e, bottom), accompanied by a broadly extended pattern of Rcs activation (Fig. 4d,f, bottom, Extended Data Fig. 9, Supplementary Videos 12– 15). In open channels, aligned cells exhibited weak Rcs activation over time (Extended Data Fig. 7c, Supplementary Video 16). Together, these results indicate that confinement geometry governs cell orientation patterns and defect distributions, thereby enabling spatial biasing of Rcs activation. While this coupling does not strictly determine the spatial pattern of Rcs activation, it provides a means to regulate where Rcs activation emerges within the chamber.

## 3 Discussion

In this study, we show that cell orientation patterns and associated topological defects contribute to mechanically induced activation of the Rcs pathway, a key component of extracellular matrix production, in *E. coli* biofilm. Specifically, Rcs activation preferentially emerges in regions influenced by topological defects in growing populations, linking defect-associated mechanical cues to the initiation of biofilm formation. Because pressure is known to be a mechanical cue for cells to induce Rcs activation [11], our findings suggest that the characteristic pressure generated by topological defects contributes to the activation of the Rcs pathway. Consistent with this interpretation, we were able to bias Rcs activation sites by designing cell orientation patterns and defect distributions using square-shaped microfluidic chambers. Notably, Rcs activation was observed both near topological defects and near chamber corners, suggesting that the spatial organization of defects together with confinement-mediated redistribution of mechanical stress may contribute to its patterning. These results indicate that cell orientation patterns and topological defects are associated with the spatial emergence of Rcs activation in confined *E. coli* populations, suggesting a possible biophysical contribution to an early stage of biofilm initiation. More broadly, our findings suggest that cell orientation patterning may help modulate the onset and spatial organization of stress-response pathways involved in matrix production, with potential implications for early biofilm development.

Although the present study and prior work on topological defects in bacterial assemblies have focused on rod-shaped species [17–21, 27], mechanical interactions associated with defects may provide insight into collective behaviors across broader bacterial morphologies. Orientational order has been observed not only in straight rods but also in curved bacteria such as *Vibrio cholerae* and *Caulobacter crescentus*, which form microdomains within populations [46]. Even coccoid and ovoidal species, such as *Staphylococcus aureus, Streptococcus pneumoniae, Enterococcus faecalis*, and *Lactococcus lactis*, exhibit transient elongation during growth and division, which may result in the cell orientation ordering [47, 48]. Thus, mechanically induced gene expression around topological defects may represent a generalizable mechanism for controlling multicellular behaviors in diverse bacteria.

An important next step is to quantitatively determine whether the pressure generated by cell–cell interactions at individual defects is sufficient to trigger Rcs activation. Approaches such as deformable microfluidic confinements [7, 8] and movable force sensors [49] have provided population-level pressure measurements, but single-cell or submicron-scale in situ quantification remains challenging. Future studies could benefit from microfabricated deformable gels [50, 51] and the mechanical properties of soft cells [52]. Furthermore, studies including the present one and others [8, 11] demonstrate that fluorescence reporters associated with the Rcs pathway can serve as indirect but spatially precise readouts of mechanical loading. Although the relationship between quantified pressure, Rcs activation, and subsequent CA production requires further investigation, integrating mechanical measurements with reporters of Rcs activation and CA production may enable single-cell-level mapping of pressure landscapes around individual topological defects.

Finally, our findings suggest that controlling bacterial orientation patterns could provide a strategy to suppress biofilm formation. Spatial control of topological defect positions has been demonstrated in mammalian cells using surface-patterned substrates [53–56]. Applying similar principles to bacterial populations could reduce mechanically induced extracellular matrix production by relocating or diluting topological defects. Because extracellular matrix protects cells from antibiotics [57], reduced Rcs activation and CA production may enhance the efficacy of conventional treatments. Additionally, weakened matrix production could mechanically destabilize early biofilms and increase susceptibility to hydrodynamic removal [58]. Thus, combining spatial control of topological defect positions with existing biofilm-suppression strategies may provide a multifaceted approach for preventing biofilm-associated infections by limiting mechanically induced matrix production at its origin.

## 4 Methods

### 4.1 Bacterial strains and pre-culture preparation

We used an engineered strain derived from *E. coli* K-12 MG1655 [11], which does not produce a typical extracellular matrix of *E. coli*, cellulose, as it has a mutation in the *bcsQ* gene of the parent K-12 strain [59, 60]. This *E. coli* strain was prepared by a general transformation method to obtain two plasmids for constitutive and inducible fluorescence proteins. Rcs activation was quantified using fast-maturating msfGFP [35], with a kanamycin resistant marker for selection, under the native promoter of *rprA*, which is in the Rcs pathway and simultaneously upregulated with the *cps* gene, which is directly involved in CA production [15]. This Rcs pathway is known to be triggered by pressure such as substrate friction, osmotic responses, and cell deformation [10, 61]. By measuring this msfGFP fluorescence intensity, Rcs activation has been quantified under pressure to *E. coli* cells [11].

For a control experiment of induced Rcs activation, this *E. coli* strain was transformed with a plasmid of pZA3R-tdTomato for constitutive fluorescence protein production. This plasmid was extracted from another *E. coli* strain using a plasmid extraction kit (QIAprep Spin Miniprep Kit, Qiagen) according to the manufacturer’s protocol. Then, the parent strain at the exponential growth phase (optical density at 600 nm (OD_600_) around 0.3) was used for transformation with CaCl_2_ [62]. Briefly, 1 mL of cultured cells was collected by centrifuging at 25 °C and 5, 000 rpm for 5 min using a centrifuge (himac CT13R, Hitachi Koki) and suspended in 300 *µ*L of 100 mM CaCl_2_ with incubation on ice for 2 min. The suspended cells were collected again with the same centrifugation step and suspended in 60 *µ*L of 100 mM CaCl_2_ by pipetting with incubation on ice for 5 min. One microliter of plasmid solution was added to the suspension. This cell suspension was incubated on ice for 15 min, then on a heat block at 42 °C for 90 sec for heat shock. Immediately after this step, the cells were incubated on ice for 2 min. One milliliter of LB (BD Biosciences) was added to the cell suspension, followed by incubation at 37 °C for 30 min. As the plasmid pZA3R-tdTomato has a gene for chloramphenicol resistance, 20 *µ*L of the transformed cell suspension was seeded and dried on a 1.5 w/v% agar plate with kanamycin (50 *µ*g/mL) and chloramphenicol (100 *µ*g/mL) and cultured at 37 °C for approximately 17 h for the subsequent positive selection. When colonies appeared on the plate, some of them were picked up and suspended in an aliquot of LB using a pipette tip and 2 *µ*L of the suspension was observed under a microscope to investigate whether the tdTomato fluorescence intensity was detected from the transformed cells (see Methods 4.2 below). A transformed strain was cultured in 1 mL of LB at 37 °C for approximately 17 h, then the cultured cell medium was stored in 10 v/v% of glycerol at −80 °C.

For imaging experiments, cell culture at the exponential growth phase was prepared as follows. A glycerol stock of this *E. coli* was streaked onto an LB plate containing 1.5 w/v% agar and incubated at 37 °C for 24 h and stored at 4 °C until used for less than three months. A part of colony was inoculated in LB with kanamycin (50 *µ*g/mL) and chloramphenicol (100 *µ*g/mL) of a 4mL culture tube (polypropylene, 2332-004S, Watson) and cultured at 200 rpm and 37 °C for 17 h using a shaker incubator (BioShaker BR-21FP, Titec). Two microliters of the pre-cultured medium was subcultured in 1 mL of LB with kanamycin (50 *µ*g/mL) and chloramphenicol (100 *µ*g/mL) and cultured at 200 rpm and 37 °C approximately for 2 h until cells entered the exponential growth phase ( OD_600_ around 0.3). This cultured medium was used for the imaging experiments, both using agar pad and microfluidics.

### 4.2 Imaging under agar pads

For overnight time-lapse imaging using an agar pad, we used a sealed system similar to the one used in [17]. A square-shaped double adhesive tape (SLF0601, Bio-Rad Laboratories) was put on a No. 1 glass coverslip (0.13–0.17 mm thick, C024601, Matsunami Glass). Four microliters of pre-culture cell suspension was put on the glass coverslip and covered with a square piece of LB agar pad (1.5 w/v% agar, approximately 6 mm thick by pouring 40 mL solution into a 9-cm diameter dish) with kanamycin (50 *µ*g/mL) and chloramphenicol (100 *µ*g/mL). A 3D-printed frame, whose size was almost identical to that of the tape, was attached to it (inner side 15 mm, outer side 23 mm). Another square-shaped double adhesive tape with a plastic cover was attached onto the 3D-printed frame, then placed on a microscope stage with incubation at 37 °C for 30 min for heat stabilization before imaging.

This agar pad device was observed using a confocal microscope (SP8, Leica) with a 100 *×* oil-immersion objective (HC PL APO C52, N.A. 1.40, 11506372, Leica) under Leica LasX operation with 400 Hz scan speed, 5-min time interval for 17 h, 0.5 *µ*m z-stack interval for 3-*µ*m image height, and 1 Airy unit for confocal pinhole. The image size was 512 *×* 512 pixels with a pixel size ≈ 0.23 *µ*m. Brightfield images were taken simultaneously with tdTomato fluorescence images. The msfGFP fluorescence images were taken using a HyD detector and a 488 nm laser (intensity 10 %) for excitation and a 500– 540 nm detection range for emission with an average of two line scans. The tdTomato fluorescence images were taken using a HyD detector and a 552 nm laser (intensity 1 %) for excitation and a 570–610 nm detection range for emission. We imaged multiple positions of each setup and repeated the same experiment using two independently prepared setups. We picked up z-planes of the stack images at the bottom and 3 *µ*m higher to quantify the cells protruding from the bottom colony. Two biologically independent cultures were used for each experiment, with one device per culture. A total of 10 imaging positions across the 2 devices were analyzed.

### 4.3 Master mold fabrication

Master molds of the microfluidic devices were developed on a 2-inch Si wafer (ID: 444, University Wafer) using a negative photoresist, SU-8. The development process has three steps. First, after wafer activation by baking it at 95 °C for 5 min and plasma treatment for 30 sec using a plasma cleaner (PDC-32G, Harrick Plasma) with atmospheric air and the high power mode, two alignment marks composed of four small squares with 10-*µ*m gaps between each were developed on a wafer. Second, after the wafer activation again next day, the first layer of cell culture chambers was developed: SU-8 2 (Nippon Kayaku) was spincoated on the wafer at 3, 000 rpm for 30 sec, baked at 65 °C for 1 min, then at 95 °C for 1 min, exposed to 350 *µ*J/cm^2^ of 365 nm LED using a mask-less aligner ( *µ*MLA with Writing Mode II, Heidelberg Instruments), then baked at 65 °C for 1 min, then at 95 °C for 1 min. Third, the second layer of cell culture chambers was developed: SU-8 3025 (Nippon Kayaku) was spincoated on the wafer at 4, 000 rpm for 30 sec, baked at 95 °C for 10 min, exposed to 350 *µ*J/cm^2^ of 365 nm LED, then baked at 65 °C for 1 min, then at 95 °C for 5 min. The patterned structures were developed in SU-8 developer (Nippon Kayaku) by gently shaking at 25 °C for 5 min, then in a fresh one for 30 sec, rinsed with isopropanol, and blown with N_2_ gas. The developed wafer was coated with a gas phase of Trichloro(3,3,4,4,5,5,6,6,7,7,8,8,8-tridecafluorooctyl)silane (Fujifilm Wako Pure Chemical) in a vacuum chamber at 25 °C for approximately 15 min. The developed structure height was profiled using a surface profiler (Dektak XT-A, Bruker).

### 4.4 Microfluidic device fabrication

PDMS microfluidic devices were fabricated using a 10 : 1 mass ratio of Sylgard 184 silicone elastomer base and curing agent (Dow, Midland, MI). These reagents were mixed well with a plastic spatula and degassed under vacuum for approximately 30 min, until no visible bubbles remained. Approximately 9.9 g of the mixture was poured onto the master mold with a 3D-printed frame (approximately 4 cm in diameter) and placed at 60 °C for approximately 17 h. Afterwards, the cured PDMS was peeled off from the master mold, and unnecessary parts of the PDMS block were cut off. The thickness of the device was approximately 5 mm. Two inlets/outlets were punched using a 1.5-mm biopsy puncher (Kai Corporation). The surfaces of the device and a No. 1 microscopy glass coverslip (0.13–0.17 mm thick, C024601, Matsunami Glass) were cleaned using a removable adhesive tape (Scotch Magic Mending Tape, 3M). The device and the glass coverslip were plasma-activated using a PDC-32G plasma cleaner with atmospheric air and the high power mode for approximately 1 min and bonded together. The glass-bonded device was put on a heater at 95 °C for approximately 10 min and stored at 25 °C until use, not longer than 24 h.

### 4.5 Imaging on microfluidics

Twenty microliters of 0.1 w/v% bovine serum albumin (BSA, Sigma-Aldrich) was taken up using a P20 micropipette, and the pipette tip was inserted into a 1.5-mm punched hole on a microfluidic device. An empty pipette tip was inserted into the other hole in the same way. The pipette tips were cut slightly above the liquid level of the medium using a pair of nippers. The device with the pipette tips was then centrifuged at 25 °C and 1, 000 *× g* for 5 min using a plate centrifuge (PlateSpin3, Kubota) and incubated at 25 °C for approximately 30 min, coating the device surface with BSA to prevent bacterial cell adhesion. One milliliter of pre-cultured medium was concentrated tenfold using a centrifuge (himac CT13R) at 25 °C and 5, 000 rpm for 10 min. A P20 micropipette tip filled with 20 *µ*L of concentrated pre-cultured cell suspension and an empty one were inserted into a 1.5-mm punched holes on a microfluidic device and cut in the same manner described above. The device with the pipette tips was then centrifuged at 25 °C and 1, 000 *× g* for 5 min using the plate centrifuge (PlateSpin3), loading cells into confiner chambers. The device was connected with a polytetrafluoroethylene tube (PTFE, inner diameter: 0.25 mm, outer diameter: 1.59 mm, 6010-35603, GL Sciences) filled with LB containing kanamycin (50 *µ*g/mL) and chloramphenicol (100 *µ*g/mL) by a syringe (20 mL, SS-20ESZ, Terumo, Tokyo, Japan) and a syringe pump (Pump 11 Elite, Harvard Apparatus). This microfluidic system was placed on a microscope stage with a stage-top incubator (STXG-WSKMX-SET, Tokai Hit), followed by incubation at 37 °C with 0.5 mL/h medium flow into the device using the syringe pump for 30 min for heat stabilization before imaging. During imaging, the medium was supplied at 0.1 mL/h into the device. When cell suspension was introduced to the device, some cells adhered to the substrate or diffused in the chamber (Supplementary Videos 3 or 4, respectively). Cells in confiner chambers of this microfluidic device were observed under the same conditions as the imaging under agar pads (see Methods 4.2) except for z-scan. Two or three biologically independent cultures were used for each experiment, with one device per culture, for the circular or square confiner device experiments, respectively.

For the circular confiner device experiments, a total of 6 imaging positions across the 2 devices were analyzed. For the square 20-*µ*m confiner device experiments with the splay configuration, a total of 4 square corners across the 2 imaging positions and 2 devices were analyzed. For the square 20-*µ*m confiner device experiments with the bend configuration, a total of 6 square corners across the 3 imaging positions and 3 devices were analyzed. For the square 20-*µ*m confiner device experiments with the mixed configuration, a total of 6 square corners across the 3 imaging positions and 2 devices were analyzed. For the square 30-*µ*m confiner device experiments, a total of 6 square corners across the 3 imaging positions and 1 device were analyzed.

### 4.6 Image analysis: Colony growth using brightfield images

All image analyses using Fiji/ImageJ were performed with version 1.54p. To quantify colony growth in monolayer confiner chambers, we analyzed brightfield images. The images were inverted, and their backgrounds were subtracted using the rolling ball method (radius = 10 pixels). A binary circular mask matching the chamber geometry was generated with the Create Mask function and inverted. Subtracted brightfield images were then multiplied by this mask to exclude chamber boundaries from the analysis. The resulting images were processed with the Find Edge function and converted into binary masks using Otsu thresholding [63] with a dark background. Colony areas were quantified with the Analyze Particles function using a size cutoff of 2 *µ*m^2^ to infinity.

### 4.7 Image analysis: Rcs stress-response activation by protruding cells

To detect and quantify protruding regions of cells from the colony bottom, we used brightfield z-scan images. Brightfield images at the colony bottom and 3*µ*m above were inverted, and their backgrounds were subtracted using the rolling ball method (radius = 5 pixels, a pixel size ≈ 0.23 *µ*m). The processed images were converted into binary masks using Otsu thresholding method [63] with a dark background, followed by six iterations of 1-pixel dilation and twelve iterations of 1-pixel erosion, and were then inverted again. This procedure generated two image series: one containing protruding regions and the other containing outside and edge regions of colonies. To remove colony edges—which often exhibited high gray values and were misidentified as protrusions—we subtracted the outside/edge series from the protrusion series. The resulting images were converted again into binary masks (Otsu thresholding, dark background), eroded twice, and dilated twice to suppress noise and bridge gaps between protruding cells. These masks were overlaid on brightfield images of the bottom colony to visualize protrusions (Fig. 1b, Supplementary Videos 1, 2). Areas of protruding regions and bottom colonies were then quantified using the Analyze Particles function with a size cutoff of 0.5 *µ*m^2^ to infinity. To normalize images across locations, the time point at which protruding regions exceeded 200 *µ*m^2^ was defined as 0 min. The non-protruding areas were defined as the regions remaining after subtracting protruding areas and the outside and edge regions of colonies. Rcs activation was defined as the total gray values of msfGFP at the bottom images.

### 4.8 Image analysis: Cell orientation and topological defects

To visualize and quantify cell orientation fields, we used brightfield images of cells in monolayer confiner devices. Time-lapse multiple images of brightfield were aligned using StackReg plugin using rigid body transformation [64]. Then, the aligned images were cropped using a square within the monolayer confiner chambers. The cropped images were analyzed using a Fiji/ImageJ plugin, OrientationJ [40]. Images were processed by the OrientationJ Analysis function using a Gaussian filtering with a local window *σ* = 5 pixels (a pixel size ≈ 0.23 *µ*m), and the resulting images were exported after color survey with orientation hue and coherency saturation on the original images. Their brightness was modified manually in the same manner for all the images. These images were processed by the OrientationJ Distribution function using the same parameters to quantify orientation fields.

To detect topological defects, we used a custom package known as Defector [18] on MATLAB with the same *σ* = 5 pixels (a pixel size ≈ 0.23 *µ*m).

### 4.9 Image analysis: Rcs stress-response activation in monolayer confiner devices

We selected chambers in which cells fully occupied the space and initially exhibited relatively low msfGFP fluorescence intensity. During preliminary observations of precultured cells, we noticed two distinct subpopulations: one with longer rod-shaped cells showing lower msfGFP fluorescence, and the other with shorter or spherical cells displaying much brighter signals (Extended Data Fig. 10). The latter cells did not grow over time, suggesting that they were dormant or dead and did not contribute to Rcs activation in the growing population. Imaging positions including such abnormal cells were detected manually and excluded from the further analysis. To normalize across different chambers, we defined the initial time point as the moment when cells first filled the chamber. Before the analysis, We defined the onset of Rcs activation (*T*_onset_) as the time point when the maximum msfGFP fluorescence increased by at least 20 units from the initial value, marking the initiation of biofilm formation.

Using these criteria, we analyzed images of growing cells in the chambers. For a fair comparison in circular colonies, we used ratios calculated by dividing the high Rcs activation pixel number by the total pixel number within the corresponding bins (Extended Data Fig. 5), where high Rcs activation pixels were defined as those with msfGFP fluorescence intensities exceeding the maximum value observed in the initial state.

### 4.10 Image analysis: Cell orientation in square confiner devices

To quantify cell orientation fields in the square confiner chambers, we followed the same method, described above (see Methods 4.8) using image data *±* 20 min after chamber filling with slight modifications. After the alignment, brightfield images of square confiner chambers were cropped with a square and rotated at −45° to fill the square field of view with the images taken by the experiments. Then, these rotated images were used in orientation analysis. For visualization, the analyzed images were rotated back at 45° to the original image angle and overlaid using the maximized saturation to clear visualization.

### 4.11 Image analysis: Splay and bend classification in square confiner devices

To classify cell orientation patterns as predominantly splay or bend, images were analyzed at time points when cells fully occupied square chambers. Orientation fields were extracted using the OrientationJ Vector Field plugin using a Gaussian filtering with a local window *σ* = 5 pixels (a pixel size ≈ 0.23 *µ*m) and a grid size of 10 pixels. Only vectors with coherency ≥ 0.1 were retained for analysis. Unit director vectors were then defined as *n*_*x*_ = cos(*θ*_*n*_) and *n*_*y*_ = sin(*θ*_*n*_), where *θ* is the local orientation angle. Local orientation modes (splay or bend) were quantified using the scalar metric (**r** · **n**)^2^, where **r** is the unit radial vector from the closed corner of each square chamber as a reference point and **n** is the local director. Three sampling points were defined near the reference points. At each point, (**r** · **n**)^2^ was calculated and classified using predefined thresholds: values ≥ 0.65 were assigned as splay, ≤ 0.35 as bend, and intermediate values as mixed. Mean values were computed for each region, and a global score was obtained by averaging across regions. The overall orientation mode was determined from the global score using the same classification thresholds.

## 5 Data and code availability

Relevant experimental data, together with the Fiji/ImageJ and MATLAB scripts used to quantify protruding cell area and spatial Rcs activation, are available via Zenodo at https://doi.org/10.5281/zenodo.19200174. Due to repository size constraints, only representative image datasets are publicly deposited. All other data supporting the findings of this study are available from the corresponding author upon reasonable request.

## Acknowledgement

We thank G. Mason and E.R. Rojas for sharing the *E. coli* strain fluorescently reporting Rcs activation. We thank Y. Wakamoto and R. Okura for helping us to use their surface profiler. We thank Y.T. Maeda and H. Salman for sharing the plasmid DNA pZA3R-tdTomato. This work is supported by KAKENHI from Japan Society for the Promotion of Science (JSPS) (No. JP22K20564, JP23K14516, and JP23KJ0327 to FY and JP24K00593 and JP19H05800 to KAT), KAKENHI for JSPS Fellows (No. JP23KJ0327 to FY), Japan Science and Technology Agency (JST) PRESTO (JPMJPR25NG to FY) and FOREST (No. JPMJFR2364 to KAT), the Nakatani Foundation Incentive Research Grant (2023S238 to FY), the JSPS Core-to-Core Program “Advanced core-to-core network for the physics of self-organizing active matter (No. JPJSCCA20230002), and the JSPS Program for Forming Japan’s Peak Research Universities (J-PEAKS) (No. JPJS0042023001).

## Author contributions

F.Y. conceived the project, conducted all the experiments, analyzed the data, designed the figures, and wrote the draft. F.Y. and K.A.T. secured funding, interpreted the data, and wrote the manuscript.

## Additional information

**Correspondence and requests for materials** should be addressed to Fumiaki Yokoyama.

## Extended Data Figure

### Supplementary Video Legends

**Extended Data Fig. 1.**
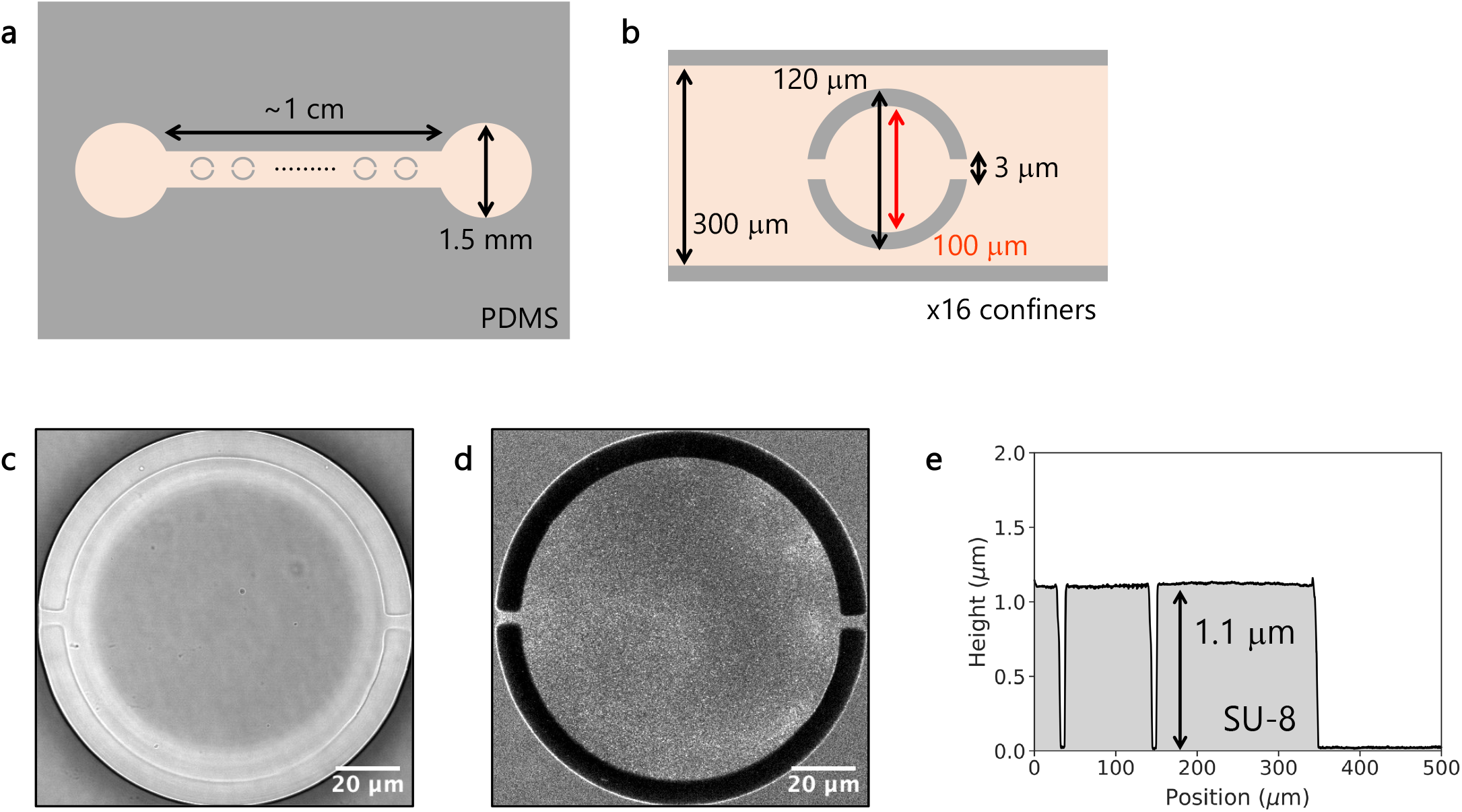
Characterization of monolayer confiner microfluidic device. **a**, A sketch of the whole device design. An inlet and an outlet are connected via a channel containing several monolayer confiner chambers. **b**, A sketch of a monolayer confiner chamber. These sketches are not to scale. **c**, A representative brightfield image of a monolayer confiner chamber attached to a glass coverslip. **d**, A representative autofluorescence image of LB flowing through a monolayer confiner chamber attached to a glass coverslip. Parts of the chamber attached to a glass substrate show no fluorescence signals. **e**, Height characterization of a monolayer confiner chamber on the SU-8-patterned Si wafer.

**Extended Data Fig. 2.**
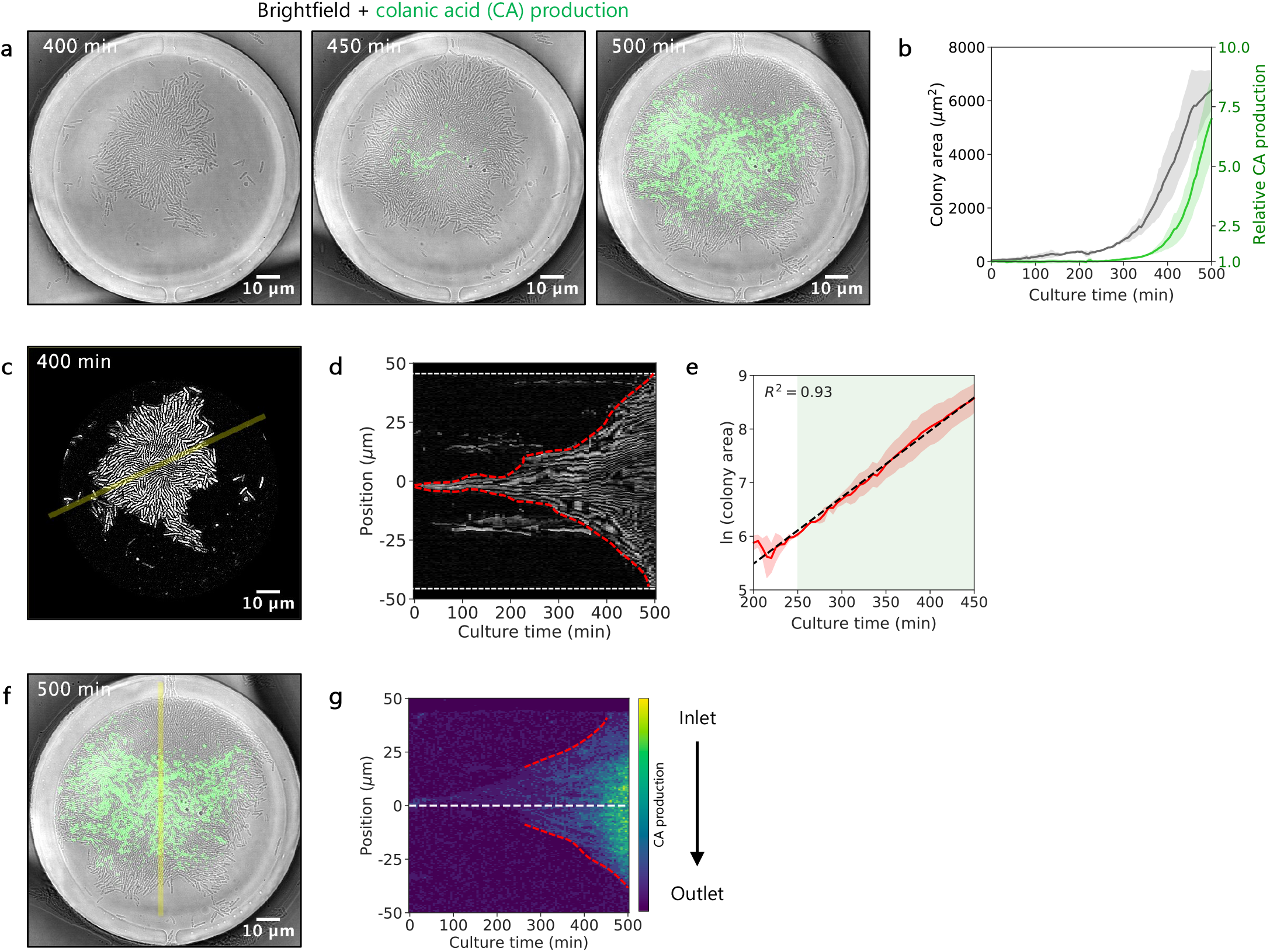
Cell growth and Rcs stress-response activation in monolayer confiner devices. **a**, Representative time-lapse images of attached *E. coli* cells growing and activating the Rcs pathway in a monolayer confiner chamber. The green color shows fluorescence intensity of msfGFP associated with Rcs activation. **b**, Time evolution of colony area and relative msfGFP fluorescence intensity to the initial value as Rcs activation. **c**, A representative brightfield image of cells in a monolayer confiner chamber after segmentation, with a yellow line for creating the kymograph in **d. d**, A kymograph of cells from the segmented brightfield images in a monolayer confiner chamber. The kymograph was made along the yellow line in **c**. The red dashed lines indicate the edge of the segmented colony. **e**, Time evolution of the natural logarithm of colony area. The black dashed line shows a linear regression fit. The green shade shows the time range of Rcs activation over 250 −450 min, as shown in **b. f**, An overlaid image of brightfield and msfGFP fluorescence associated with Rcs activation of attached cells, with a yellow line for creating the kymograph in **g. g**, A kymograph of msfGFP fluorescence intensity in a monolayer confiner chamber. The kymograph was made along the yellow line in **f**. The msfGFP fluorescence intensity reporting Rcs activation is shown using the viridis color scale. The black line shows the direction of the medium flow from the inlet to the outlet through the monolayer confiner chamber. The white dashed line indicates the center position of the kymograph. The red dashed lines indicate the edge of the Rcs activation areas. Each line plot shows the mean with standard errors (2 positions across 2 devices were used).

**Extended Data Fig. 3.**
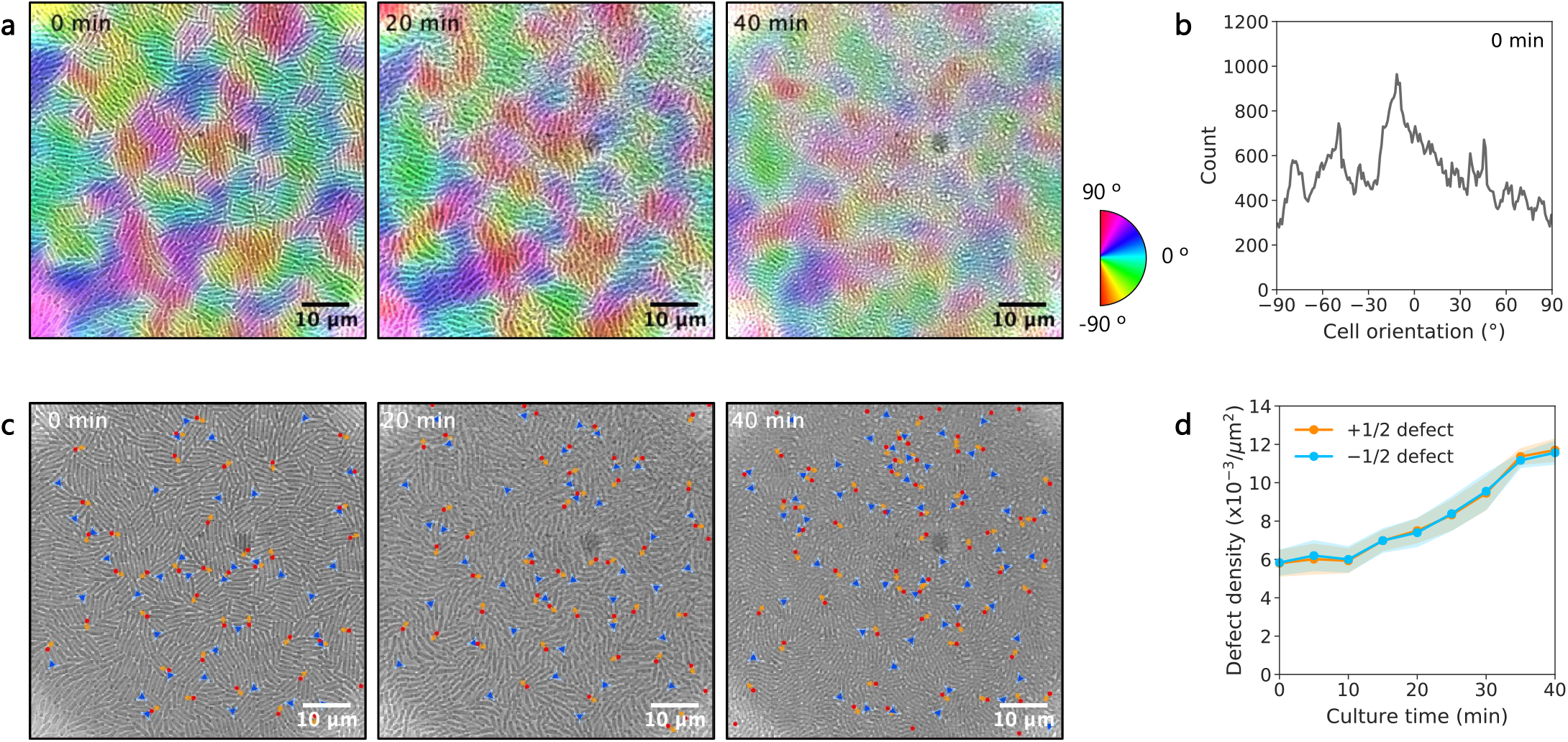
Cell orientation and topological defects in monolayer confiner devices. **a**, Representative time-lapse images of brightfield with color codes. **b**, A representative distribution of cell orientation at 0 min. **c**, Representative time-lapse images of brightfield with *±* 1/2 topological defects. The red circles and blue triangles show +1/2 and −1/2 topological defects, respectively. **d**, Density of *±* 1/2 topological defects over time. Each line plot except for **b** shows the mean with standard errors (6 positions across 2 devices were used).

**Extended Data Fig. 4.**
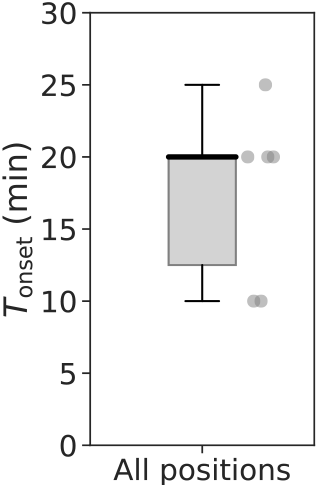
Timing of high Rcs activation. A boxplot of timing at the first time point at *T*_onset_ in monolayer confiner chambers. The black line within the boxes represents the median, the lower and upper edges of the boxes represent the 25 and 75 percentiles of the data, respectively, and the error bar represents the maximum and minimum values of the box plots (6 positions across 2 devices were used).

**Extended Data Fig. 5.**
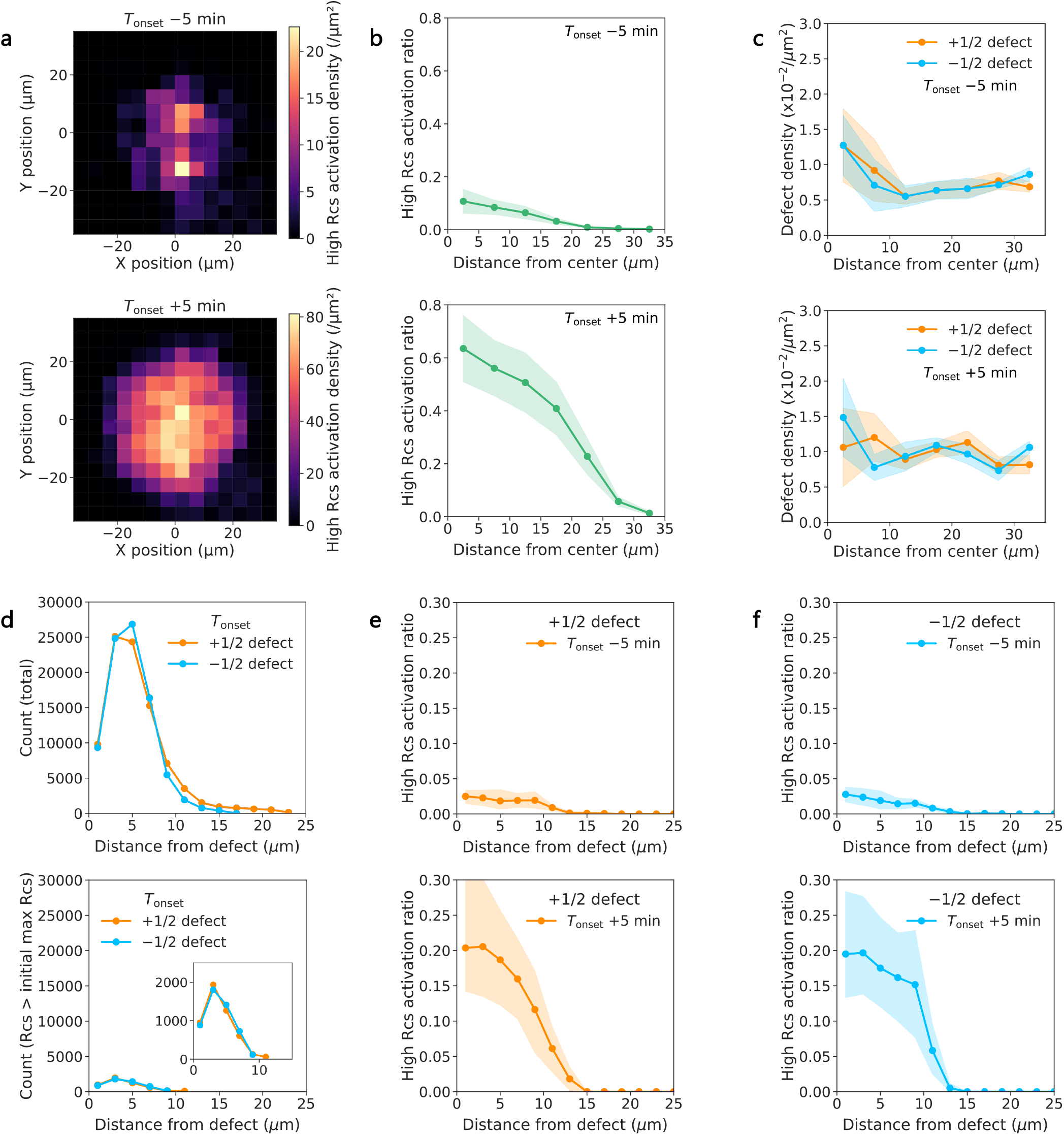
Histogram of total pixel number and high Rcs activation pixels. **a**, 2D distribution of high Rcs activation pixel densities over colony at 5 min before and after *T*_onset_ (top and bottom, respectively) (see Fig. 3c at *T*_onset_). **b**, 1D distribution of high Rcs activation pixel ratios as a function of the distance from the colony center at 5 min before and after *T*_onset_ (top and bottom, respectively) (see Fig. 3d at *T*_onset_). The ratio of high Rcs activation pixels was calculated by dividing the number of high Rcs activation pixels by the total pixel number. High Rcs activation pixels were defined as those with msfGFP fluorescence intensities exceeding the maximum value observed in the initial state, when cells fully occupied the chamber and maintained their orientations at the subsequent time point. **c**, Density of *±*1/2 topological defects as a function of the distance from the colony center at 5 min before and after *T*_onset_ (top and bottom, respectively) (see Fig. 3e at *T*_onset_). **d**, A representative 1D distribution of total and high Rcs activation pixel number (top and bottom, respectively) as a function of the distance from +1/2 and −1/2 topological defects at *T*_onset_. The inset in the bottom panel shows a close-up. **e,f**, High Rcs activation pixel ratios as a function of the distance from +1/2 (**e**) and −1/2 (**f**) topological defects at 5 min before and after *T*_onset_ (top and bottom, respectively) (see Fig. 3f at *T*_onset_). Each line plot shows the mean with standard errors (6 positions across 2 devices were used).

**Extended Data Fig. 6.**
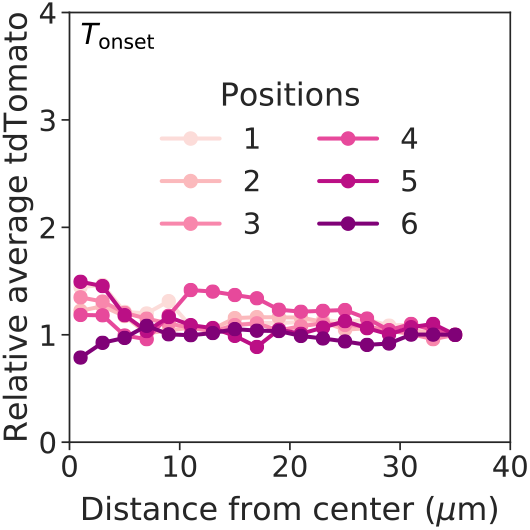
Distribution of a non-inducible fluorescence protein in monolayer confiner devices. Relative values of constitutive tdTomato fluorescence intensity over colonies at *T*_onset_. The relative values were normalized with those at the edge of colony (35 *µ*m from colony center). This line shows data from different fields of view (6 positions across 2 devices were used).

**Extended Data Fig. 7.**
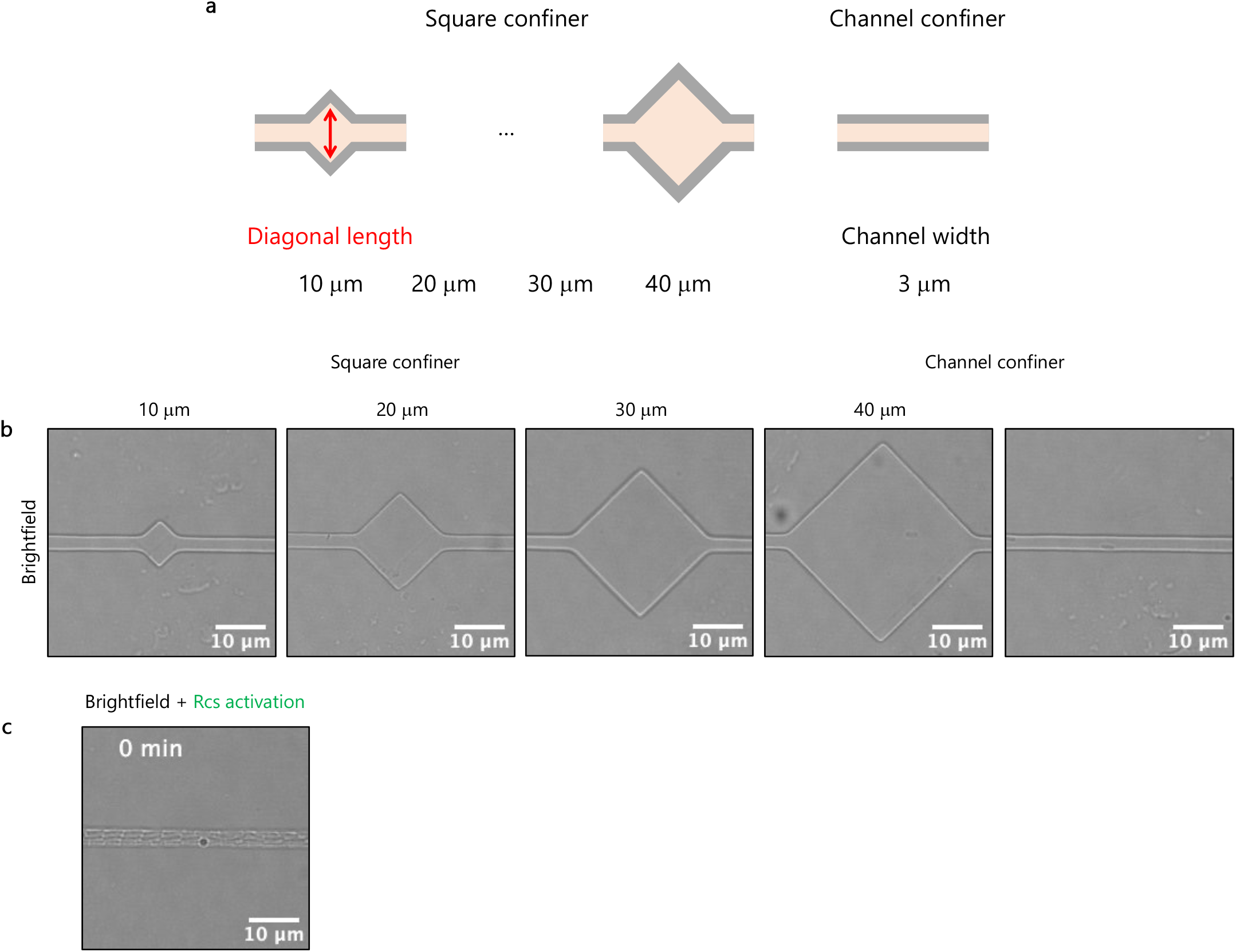
Rcs stress-response activation in square confiner devices. **a**, Sketches of square and channel confiners. These sketches are not to scale. **b**, Brightfield images of square and channel confiners. **c**, Overlaid images of brightfield and msfGFP fluorescence intensity of cells growing in channel confiner. The green color indicates msfGFP fluorescence intensity associated with Rcs activation production.

**Extended Data Fig. 8.**
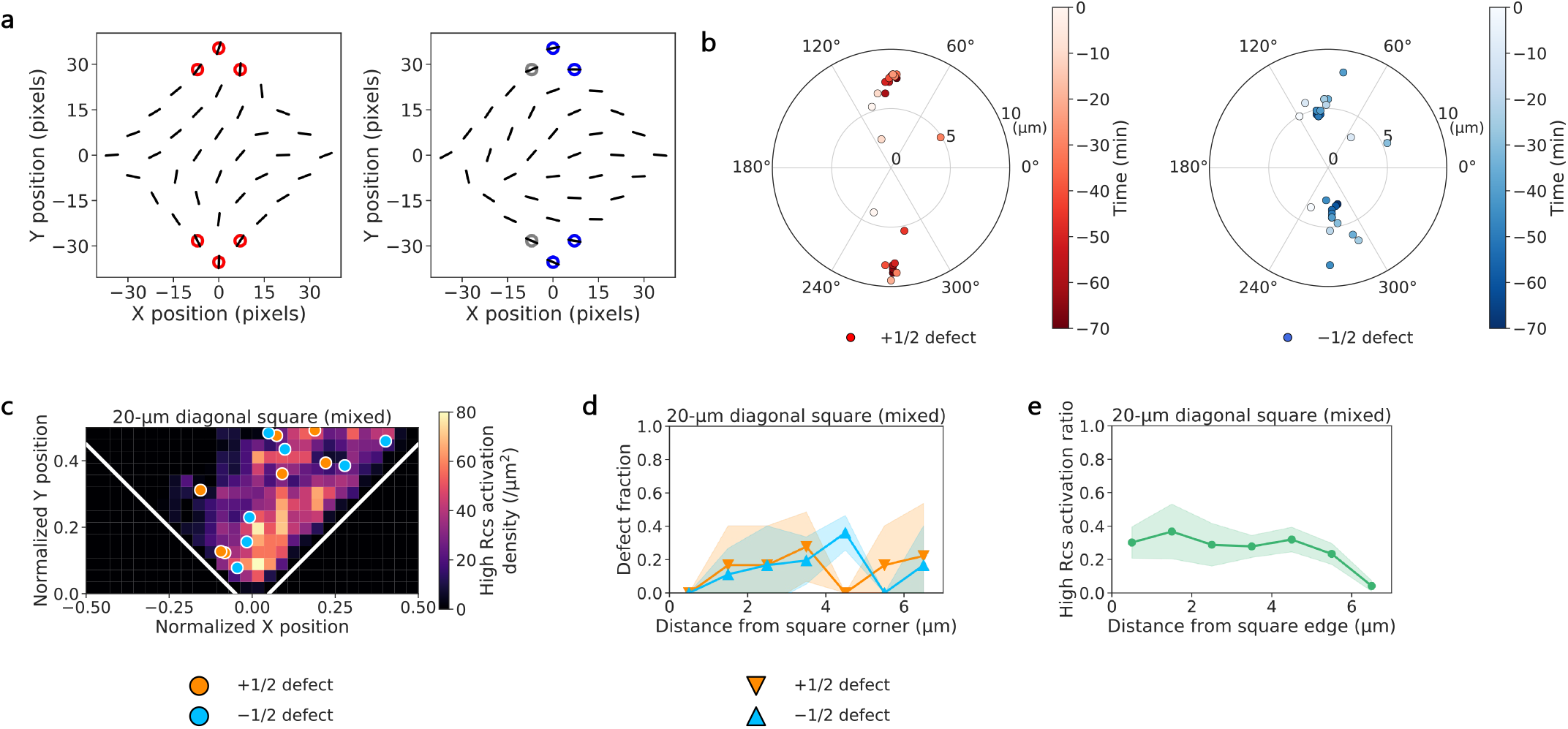
Cell orientation patterns and topological defects in square confiner devices. **a**, Representative splay and bend classifications. Orientation vectors of cells in square confiner devices with diagonals of 20 *µ*m are shown as black bars. Red, blue, and gray circles denote splay, bend, and mixed patterns, respectively, at the closed corners of the square chambers. **b**, A representative time evolution of +1/2 (left) and −1/2 (right) topological defects over 70 min before *T*_onset_ in the square confiner with diagonal lengths of 20 *µ*m. The straight channel spans the square chamber from 0° to 180°, with closed corners located at 90° and 270°. **c**, Spatial distribution of representative topological defects and high Rcs activation pixel densities in square confiner devices with diagonals of 20 *µ*m exhibiting mixed cell orientation at *T*_onset_. X and Y positions are normalized by the diagonal length of the square chambers. White lines delineate the edges of the square chambers. **d**, Mean number fraction of *±* 1/2 topological defects as a function of the distance from the corner in square confiner devices with diagonals of 20 *µ*m exhibiting mixed cell orientation at *T*_onset_. **e**, Distribution of high Rcs activation pixel ratios as a function of the distance from the edge in square confiner devices with diagonals of 20 *µ*m exhibiting mixed cell orientation at *T*_onset_. Each line plot shows the mean with standard errors (≥ 2 positions across 3 devices were used)

**Extended Data Fig. 9.**
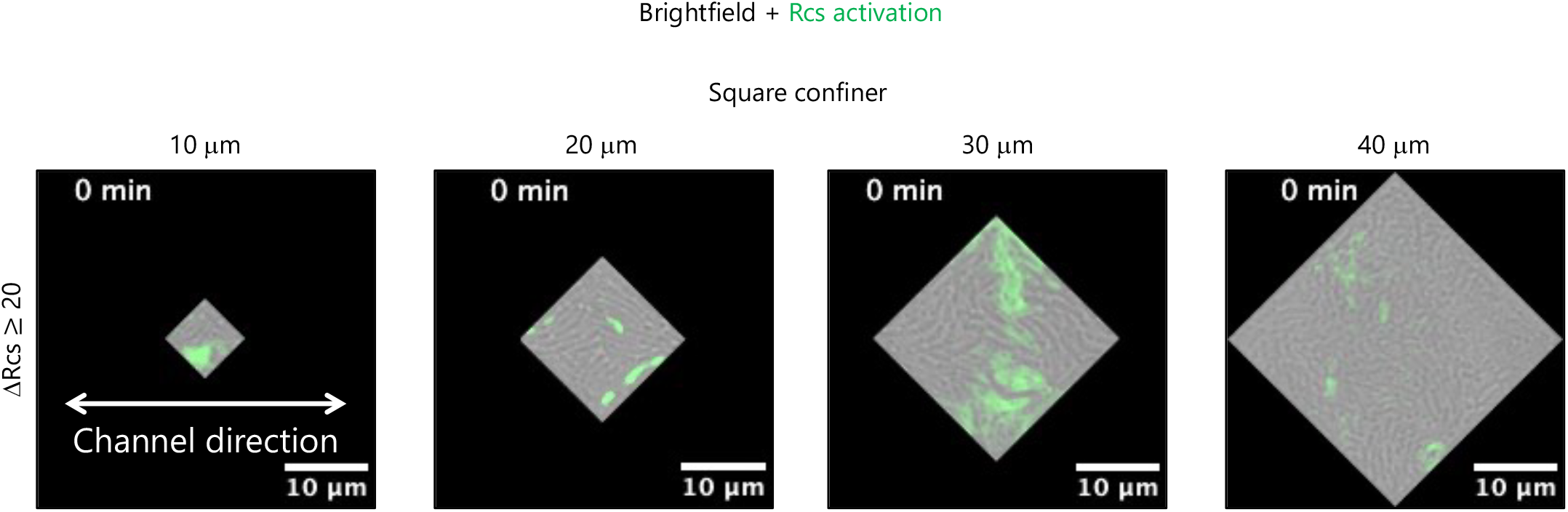
Rcs stress-response activation in square confiner chambers at *T*_onset_. **a**, Overlaid images of cropped brightfield and msfGFP fluorescence intensity of cells growing in square confiner chambers at *T*_onset_. The green color indicates msfGFP fluorescence intensity associated with Rcs activation. Channel inlets/outlets were extended along the white arrow from the square chambers (Extended Data Fig. 7).

**Extended Data Fig. 10.**
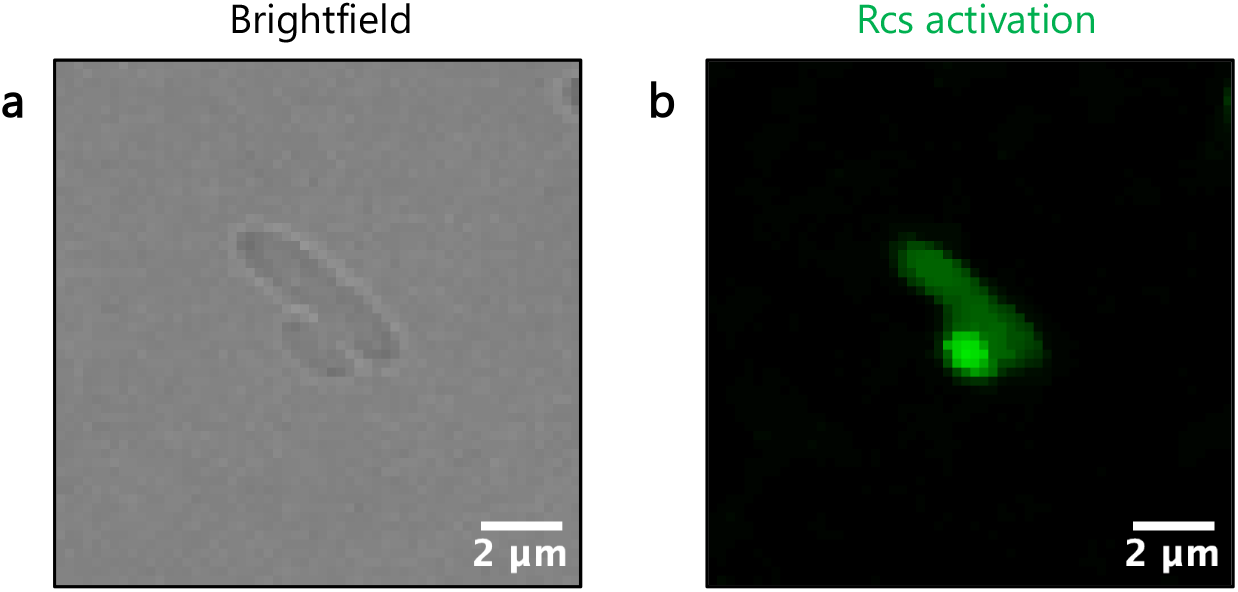
Abnormally high fluorescence intensity of round cells. **a,b**, A representative bright-field (**a**) and msfGFP fluorescence (**b**) image of normal rod-shaped and abnormal round cells.

**Supplementary Video 1**|**Cell protrusion from bottom colonies**. A representative brightfield movie of growing colonies (left) and the corresponding movie overlaid with magenta-colored protruding areas (right) of cells confined under an agar pad at a focal plane 3*µ*m above the bottom colony.

**Supplementary Video 2**|**Cell protrusion from bottom colonies and Rcs stress-response activation**. A representative overlaid movie of brightfield and msfGFP fluorescence associated with Rcs activation (green) of cells confined under an agar pad with colored protruding areas (magenta) at the bottom colony.

**Supplementary Video 3**|**Growth of adhering cells in a monolayer confiner device**. A representative overlaid movie of brightfield and msfGFP fluorescence associated with Rcs activation of cells adhering and growing in a monolayer confiner device. Time 0 min stands for the time point when the observation was started.

**Supplementary Video 4**|**Cell diffusion and growth in a monolayer confiner device**. A representative overlaid video of brightfield and msfGFP fluorescence images of cells diffusing and growing in a monolayer confiner device. Time 0 min stands for the time point when cells fully occupied the chamber surface.

**Supplementary Video 5**|**Orientation pattern of growing cells in a monolayer confiner device**. A representative movie of brightfield with orientational color codes of a cell population growing in a monolayer confiner device. The orientational color codes are the same as Fig. 2c.

**Supplementary Video 6**|**Topological defects in cell populations in a monolayer confiner device**. A representative movie of brightfield with *±*1/2 topological defects of a cell population growing in a monolayer confiner device. The red circles and blue triangles show +1/2 and −1/2 topological defects, respectively.

**Supplementary Video 7**|**Topological defects and Rcs stress-response activation in a monolayer confiner device**. A representative overlaid movie of the msfGFP fluorescence intensity of Rcs activation and *±* 1/2 topological defects of a cell population growing in a monolayer confiner device. The msfGFP fluorescence intensity reporting Rcs activation is shown using the viridis color scale. The red circles and blue triangles show +1/2 and −1/2 topological defects, respectively.

**Supplementary Video 8**|**Orientation pattern in a square confiner chamber with diagonal lengths of** 10 *µ*m **around** *T*_onset_. A representative movie of brightfield with orientational color codes of cells growing in a square confiner chamber with a 10-*µ*m diagonal around *T*_onset_. The orientational color codes are the same as Fig. 4a.

**Supplementary Video 9**|**Orientation pattern in a square confiner chamber with diagonal lengths of** 20 *µ*m **around** *T*_onset_. A representative movie of brightfield with orientational color codes of cells growing in a square confiner chamber with a 20-*µ*m diagonal around *T*_onset_. The orientational color codes are the same as Fig. 4a.

**Supplementary Video 10**|**Orientation pattern in a square confiner chamber with diagonal lengths of** 30 *µ*m **around** *T*_onset_. A representative movie of brightfield with orientational color codes of cells growing in a square confiner chamber with a 30-*µ*m diagonal around *T*_onset_. The orientational color codes are the same as Fig. 4a.

**Supplementary Video 11**|**Orientation pattern in a square confiner chamber with diagonal lengths of** 40 *µ*m **around** *T*_onset_. A representative movie of brightfield with orientational color codes of cells growing in a square confiner chamber with a 40-*µ*m diagonal around *T*_onset_. The orientational color codes are the same as Fig. 4a.

**Supplementary Video 12**|**Rcs stress-response activation in a square confiner chamber with diagonal lengths of** 10 *µ*m **around** *T*_onset_. A representative overlaid movie of brightfield and msfGFP fluorescence associated with Rcs activation of cells growing in a square confiner chamber with a 10-*µ*m diagonal around *T*_onset_.

**Supplementary Video 13**|**Rcs stress-response activation in a square confiner chamber with diagonal lengths of** 20 *µ*m **around** *T*_onset_. A representative overlaid movie of brightfield and msfGFP fluorescence associated with Rcs activation of cells growing in a square confiner chamber with a 20-*µ*m diagonal around *T*_onset_.

**Supplementary Video 14**|**Rcs stress-response activation in a square confiner chamber with diagonal lengths of** 30 *µ*m **around** *T*_onset_. A representative overlaid movie of brightfield and msfGFP fluorescence associated with Rcs activation of cells growing in a square confiner chamber with a 30-*µ*m diagonal around *T*_onset_.

**Supplementary Video 15**|**Rcs stress-response activation in a square confiner chamber with diagonal lengths of** 40 *µ*m **around** *T*_onset_. A representative overlaid movie of brightfield and msfGFP fluorescence associated with Rcs activation of cells growing in a square confiner chamber with a 40-*µ*m diagonal around *T*_onset_.

**Supplementary Video 16**|**Cell growth and Rcs stress-response activation in a channel confiner device**. A representative overlaid movie of brightfield and msfGFP fluorescence associated with Rcs activation of cells growing in a channel confiner device.

## References

[1] Cornwallis, C.K., Svensson-Coelho, M., Lindh, M., Li, Q., Stábile, F., Hansson, L.-A., Rengefors, K.: Single-cell adaptations shape evolutionary transitions to multicellularity in green algae. Nature Ecology & Evolution 7(6), 889–902 (2023) 10.1038/s41559-023-02044-6

[2] Flemming, H.-C., Wingender, J.: The biofilm matrix. Nature Reviews Microbiology 8(9), 623–633 (2010) 10.1038/nrmicro2415

[3] Stewart, P.S., Franklin, M.J.: Physiological heterogeneity in biofilms. Nature Reviews Microbiology 6(3), 199–210 (2008) 10.1038/nrmicro1838

[4] Dufrêne, Y.F., Persat, A.: Mechanomicrobiology: how bacteria sense and respond to forces. Nature Reviews Microbiology 18(4), 227–240 (2020) 10.1038/s41579-019-0314-2

[5] Harper, C.E., Zhang, W., Lee, J., Shin, J.-H., Keller, M.R., Wijngaarden, E.v., Chou, E., Wang, Z., Dörr, T., Chen, P., Hernandez, C.J.: Mechanical stimuli activate gene expression via a cell envelope stress sensing pathway. Scientific Reports 13(1), 13979 (2023) 10.1038/s41598-023-40897-w

[6] Ni, L., Huang, Y., Huang, Y., Yu, Y., Xiong, J., Wen, H., Xiao, W., Liang, H., Jin, F.: Pressorum sensing: Growth-induced compression activates cAMP signaling in pseudomonas aeruginosa. bioRxiv (2024) 10.1101/2024.07.08.602437

[7] Chu, E.K., Kilic, O., Cho, H., Groisman, A., Levchenko, A.: Self-induced mechanical stress can trigger biofilm formation in uropathogenic escherichia coli. Nature Communications 9(1), 4087 (2018) 10.1038/s41467-018-06552-z

[8] Blanc, L.L., Alric, B., Rollin, R., Xénard, L., Finn, L.R., Goussard, S., Mazenq, L., Ingersoll, M.A., Piel, M., Tinevez, J.-Y., Delarue, M., Duménil, G., Bonazzi, D.: Bacterial growth under confinement requires transcriptional adaptation to resist metabolite-induced turgor pressure build-up. bioRxiv (2024) 10.1101/2024.09.20.614086

[9] Majdalani, N., Gottesman, S.: THE RCS PHOSPHORELAY: A complex signal transduction system. Microbiology 59(1), 379–405 (2005) 10.1146/annurev.micro.59.050405.101230

[10] Kimkes, T.E.P., Heinemann, M.: How bacteria recognise and respond to surface contact. FEMS Microbiology Reviews 44(1), 106–122 (2020) 10.1093/femsre/fuz029

[11] Mason, G., Footer, M.J., Rojas, E.R.: Mechanosensation induces persistent bacterial growth during bacteriophage predation. mBio 14(6), e0276622 (2023) 10.1128/mbio.02766-22

[12] Ren, G., Wang, Z., Li, Y., Hu, X., Wang, X.: Effects of lipopolysaccharide core sugar deficiency on colanic acid biosynthesis in escherichia coli. Journal of Bacteriology 198(11), 1576–1584 (2016) 10.1128/jb.00094-16

[13] Danese, P.N., Pratt, L.A., Kolter, R.: Exopolysaccharide production is required for development of escherichia coli k-12 biofilm architecture. Journal of Bacteriology 182(12), 3593–3596 (2000) 10.1128/jb.182.12.3593-3596.2000

[14] Laubacher, M.E., Ades, S.E.: The rcs phosphorelay is a cell envelope stress response activated by peptidoglycan stress and contributes to intrinsic antibiotic resistance. Journal of Bacteriology 190(6), 2065–2074 (2008) 10.1128/jb.01740-07

[15] Wall, E., Majdalani, N., Gottesman, S.: The complex rcs regulatory cascade. Annual Review of Microbiology 72(1), 111–139 (2018) 10.1146/annurev-micro-090817-062640

[16] Langeslay, B., Juarez, G.: Microdomains and stress distributions in bacterial monolayers on curved interfaces. Soft Matter 19(20), 3605–3613 (2023) 10.1039/d2sm01498j

[17] Shimaya, T., Takeuchi, K.A.: Tilt-induced polar order and topological defects in growing bacterial populations. PNAS Nexus 1(5), pgac269 (2022) 10.1093/pnasnexus/pgac2692106.10954

[18] Meacock, O.J., Doostmohammadi, A., Foster, K.R., Yeomans, J.M., Durham, W.M.: Bacteria solve the problem of crowding by moving slowly. Nature Physics 17(2), 205–210 (2020) 10.1038/s41567-020-01070-6

[19] Copenhagen, K., Alert, R., Wingreen, N.S., Shaevitz, J.W.: Topological defects promote layer formation in Myxococcus xanthus colonies. Nature Physics 17(2), 211–215 (2020) 10.1038/s41567-020-01056-4

[20] Han, E., Fei, C., Alert, R., Copenhagen, K., Koch, M.D., Wingreen, N.S., Shaevitz, J.W.: Local polar order controls mechanical stress and triggers layer formation in myxococcus xanthus colonies. Nature Communications 16(1), 952 (2025) 10.1038/s41467-024-55806-6

[21] Prasad, M., Obana, N., Lin, S.-Z., Zhao, S., Sakai, K., Blanch-Mercader, C., Prost, J., Nomura, N., Rupprecht, J.-F., Fattaccioli, J., Utada, A.S.: Alcanivorax borkumensis biofilms enhance oil degradation by interfacial tubulation. Science 381(6659), 748–753 (2023) 10.1126/science.adf3345

[22] Grant, M.A.A., Wacłlaw, B., Allen, R.J., Cicuta, P.: The role of mechanical forces in the planar-to-bulk transition in growing escherichia coli microcolonies. Journal of The Royal Society Interface 11(97), 20140400 (2014) 10.1098/rsif.2014.0400

[23] Duvernoy, M.-C., Mora, T., Ardré, M., Croquette, V., Bensimon, D., Quilliet, C., Ghigo, J.-M., Balland, M., Beloin, C., Lecuyer, S., Desprat, N.: Asymmetric adhesion of rod-shaped bacteria controls microcolony morphogenesis. Nature Communications 9(1), 1120 (2018) 10.1038/s41467-018-03446-y

[24] Beroz, F., Yan, J., Meir, Y., Sabass, B., Stone, H.A., Bassler, B.L., Wingreen, N.S.: Verticalization of bacterial biofilms. Nature Physics 14(9), 954–960 (2018) 10.1038/s41567-018-0170-4

[25] You, Z., Pearce, D.J.G., Sengupta, A., Giomi, L.: Monoto multilayer transition in growing bacterial colonies. Physical Review Letters 123(17), 178001 (2019) 10.1103/physrevlett.123.178001

[26] Warren, M.R., Sun, H., Yan, Y., Cremer, J., Li, B., Hwa, T.: Spatiotemporal establishment of dense bacterial colonies growing on hard agar. eLife 8, e41093 (2019) 10.7554/elife.41093

[27] Yaman, Y.I., Demir, E., Vetter, R., Kocabas, A.: Emergence of active nematics in chaining bacterial biofilms. Nature Communications 10(1), 2285 (2019) 10.1038/s41467-019-10311-z1811.12076

[28] Geisel, S., Secchi, E., Vermant, J.: The role of surface adhesion on the macroscopic wrinkling of biofilms. eLife 11, e76027 (2022) 10.7554/elife.76027

[29] Nijjer, J., Li, C., Zhang, Q., Lu, H., Zhang, S., Yan, J.: Mechanical forces drive a reorientation cascade leading to biofilm self-patterning. Nature Communications 12(1), 6632 (2021) 10.1038/s41467-021-26869-6

[30] Asally, M., Kittisopikul, M., Rúe, P., Du, Y., Hu, Z., Çağatay, T., Robinson, A.B., Lu, H., Garcia-Ojalvo, J., Süel, G.M.: Localized cell death focuses mechanical forces during 3d patterning in a biofilm. Proceedings of the National Academy of Sciences 109(46), 18891–18896 (2012) 10.1073/pnas.1212429109

[31] Peng, C., Turiv, T., Guo, Y., Wei, Q.-H., Lavrentovich, O.D.: Command of active matter by topological defects and patterns. Science 354(6314), 882–885 (2016) 10.1126/science.aah69361611.06286

[32] Genkin, M.M., Sokolov, A., Lavrentovich, O.D., Aranson, I.S.: Topological defects in a living nematic ensnare swimming bacteria. Physical Review X 7(1), 011029 (2017) 10.1103/physrevx.7.011029

[33] Dell’Arciprete, D., Blow, M.L., Brown, A.T., Farrell, F.D.C., Lintuvuori, J.S., McVey, A.F., Marenduzzo, D., Poon, W.C.K.: A growing bacterial colony in two dimensions as an active nematic. Nature Communications 9(1), 4190 (2018) 10.1038/s41467-018-06370-3

[34] Doostmohammadi, A., Thampi, S.P., Yeomans, J.M.: Defect-mediated morphologies in growing cell colonies. Physical Review Letters 117(4), 048102 (2016) 10.1103/physrevlett.117.0481021601.04489

[35] Pédelacq, J.-D., Cabantous, S., Tran, T., Terwilliger, T.C., Waldo, G.S.: Engineering and characterization of a superfolder green fluorescent protein. Nature Biotechnology 24(1), 79–88 (2005) 10.1038/nbt1172

[36] Saw, T.B., Doostmohammadi, A., Nier, V., Kocgozlu, L., Thampi, S., Toyama, Y., Marcq, P., Lim, C.T., Yeomans, J.M., Ladoux, B.: Topological defects in epithelia govern cell death and extrusion. Nature 544(7649), 212–216 (2017) 10.1038/nature21718

[37] Kunoh, T., Morinaga, K., Sugimoto, S., Miyazaki, S., Toyofuku, M., Iwasaki, K., Nomura, N., Utada, A.S.: Polyfunctional nanofibril appendages mediate attachment, filamentation, and filament adaptability in leptothrix cholodnii. ACS Nano 14(5), 5288–5297 (2019) 10.1021/acsnano.9b04663

[38] Wong, J.P.H., Fischer-Stettler, M., Zeeman, S.C., Battin, T.J., Persat, A.: Fluid flow structures gut microbiota biofilm communities by distributing public goods. Proceedings of the National Academy of Sciences 120(25), e2217577120 (2023) 10.1073/pnas.2217577120

[39] Jähne, B.: Spatio-temporal image processing: Theory and scientific applications. Springer (1993)

[40] Püspöki, Z., Storath, M., Sage, D., Unser, M.: Focus on bio-image informatics. Advances in Anatomy, Embryology and Cell Biology 219, 69–93 (2016) 10.1007/978-3-319-28549-8_3

[41] You, Z., Pearce, D.J., Giomi, L.: Confinement-induced self-organization in growing bacterial colonies. Science Advances 7(4), eabc8685 (2020) 10.1126/sciadv.abc8685

[42] Doostmohammadi, A., Ladoux, B.: Physics of liquid crystals in cell biology. Trends in Cell Biology 32(2), 140–150 (2022) 10.1016/j.tcb.2021.09.012

[43] Cho, H., Jönsson, H., Campbell, K., Melke, P., Williams, J.W., Jedynak, B., Stevens, A.M., Groisman, A., Levchenko, A.: Self-organization in high-density bacterial colonies: Efficient crowd control. PLoS Biology 5(11), e302 (2007) 10.1371/journal.pbio.0050302

[44] Sheats, J., Sclavi, B., Lagomarsino, M.C., Cicuta, P., Dorfman, K.D.: Role of growth rate on the orientational alignment of escherichia coli in a slit. Royal Society Open Science 4(6), 170463 (2017) 10.1098/rsos.170463

[45] Volfson, D., Cookson, S., Hasty, J., Tsimring, L.S.: Biomechanical ordering of dense cell populations. Proceedings of the National Academy of Sciences 105(40), 15346–15351 (2008) 10.1073/pnas.0706805105

[46] Kuroda, Y., Kawasaki, T., Menzel, A.M.: Effects of curvature on growing films of microorganisms. Biophysical Journal 124(10), 1609–1617 (2025) 10.1016/j.bpj.2025.04.003

[47] Monteiro, J.M., Fernandes, P.B., Vaz, F., Pereira, A.R., Tavares, A.C., Ferreira, M.T., Pereira, P.M., Veiga, H., Kuru, E., VanNieuwenhze, M.S., Brun, Y.V., Filipe, S.R., Pinho, M.G.: Cell shape dynamics during the staphylococcal cell cycle. Nature Communications 6(1), 8055 (2015) 10.1038/ncomms9055

[48] Wheeler, R., Mesnage, S., Boneca, I.G., Hobbs, J.K., Foster, S.J.: Super-resolution microscopy reveals cell wall dynamics and peptidoglycan architecture in ovococcal bacteria. Molecular Microbiology 82(5), 1096–1109 (2011) 10.1111/j.1365-2958.2011.07871.x

[49] Douarche, C., Allain, J.-M., Raspaud, E.: Bacillus subtilis bacteria generate an internal mechanical force within a biofilm. Biophysical Journal 109(10), 2195–2202 (2015) 10.1016/j.bpj.2015.10.004

[50] Vian, A., Pochitaloff, M., Yen, S.-T., Kim, S., Pollock, J., Liu, Y., Sletten, E.M., Campás, O.: In situ quantification of osmotic pressure within living embryonic tissues. Nature Communications 14(1), 7023 (2023) 10.1038/s41467-023-42024-9

[51] Pirnat, G., Marinčič, M., Ravnik, M., Humar, M.: Quantifying local stiffness and forces in soft biological tissues using droplet optical microcavities. Proceedings of the National Academy of Sciences 121(4), e2314884121 (2024) 10.1073/pnas.2314884121

[52] Turnbull, L., Toyofuku, M., Hynen, A.L., Kurosawa, M., Pessi, G., Petty, N.K., Osvath, S.R., Cárcamo-Oyarce, G., Gloag, E.S., Shimoni, R., Omasits, U., Ito, S., Yap, X., Monahan, L.G., Cavaliere, R., Ahrens, C.H., Charles, I.G., Nomura, N., Eberl, L., Whitchurch, C.B.: Explosive cell lysis as a mechanism for the biogenesis of bacterial membrane vesicles and biofilms. Nature Communications 7(1), 11220 (2016) 10.1038/ncomms11220

[53] Zhao, Z., Li, H., Yao, Y., Zhao, Y., Serra, F., Kawaguchi, K., Zhang, H., Sano, M.: Integer topological defects offer a methodology to quantify and classify active cell monolayers. Nature Communications 16(1), 2452 (2025) 10.1038/s41467-025-57783-w

[54] Zhao, Z., Yao, Y., Li, H., Zhao, Y., Wang, Y., Zhang, H., Chaté, H., Sano, M.: Integer topological defects reveal antisymmetric forces in active nematics. Physical Review Letters 133(26), 268301 (2024) 10.1103/physrevlett.133.268301

[55] Turiv, T., Krieger, J., Babakhanova, G., Yu, H., Shiyanovskii, S.V., Wei, Q.-H., Kim, M.-H., Lavrentovich, O.D.: Topology control of human fibroblast cells monolayer by liquid crystal elastomer. Science Advances 6(20), eaaz6485 (2020) 10.1126/sciadv.aaz6485

[56] Luo, Y., Gu, M., Park, M., Fang, X., Kwon, Y., Urueña, J.M., Alaniz, J.R.d., Helgeson, M.E., Marchetti, M.C., Valentine, M.T.: Molecular-scale substrate anisotropy and crowding drive long-range nematic order of cell monolayers. Journal of the Royal Society Interface 20(204), 20230160 (2023) 10.1098/rsif.2023.0160

[57] Ciofu, O., Moser, C., Jensen, P.O., Høiby, N.: Tolerance and resistance of microbial biofilms. Nature Reviews Microbiology 20(10), 621–635 (2022) 10.1038/s41579-022-00682-4

[58] Ohmura, T., Skinner, D.J., Neuhaus, K., Choi, G.P.T., Dunkel, J., Drescher, K.: In vivo microrheology reveals local elastic and plastic responses inside three-dimensional bacterial biofilms. Advanced Materials 36(29), e2314059 (2024) 10.1002/adma.202314059

[59] Zogaj, X., Nimtz, M., Rohde, M., Bokranz, W., Römling, U.: The multicellular morphotypes of salmonella typhimurium and escherichia coli produce cellulose as the second component of the extracellular matrix. Molecular Microbiology 39(6), 1452–1463 (2001) 10.1046/j.1365-2958.2001.02337.x

[60] Re, S.D., Ghigo, J.-M.: A CsgD-independent pathway for cellulose production and biofilm formation in escherichia coli. Journal of Bacteriology 188(8), 3073–3087 (2006) 10.1128/jb.188.8.3073-3087.2006

[61] Zietek, M., Miguel, A., Khusainov, I., Shi, H., Asmar, A.T., Ram, S., Wartel, M., Sueki, A., Schorb, M., Goulian, M., Collet, J.-F., Beck, M., Huang, K.C., Typas, A.: Bacterial cell widening alters periplasmic size and activates envelope stress responses. The EMBO Journal 44(20), 5816–5833 (2022) 10.1101/2022.07.26.501644

[62] Cohen, S.N., Chang, A.C.Y., Hsu, L.: Nonchromosomal antibiotic resistance in bacteria: Genetic transformation of escherichia coli by r-factor DNA*. Proceedings of the National Academy of Sciences 69(8), 2110–2114 (1972) 10.1073/pnas.69.8.2110

[63] Otsu, N.: A threshold selection method from gray-level histograms. IEEE Transactions on Systems, Man, and Cybernetics 9(1), 62–66 (1979) 10.1109/tsmc.1979.4310076

[64] Thevenaz, P., Ruttimann, U.E., Unser, M.: A pyramid approach to subpixel registration based on intensity. IEEE Transactions on Image Processing 7(1), 27–41 (1998) 10.1109/83.650848

